# Spatial organization of the 3D genome encodes gene co-expression programs in single cells

**DOI:** 10.1101/2022.10.26.513917

**Authors:** Peng Dong, Shu Zhang, Liangqi Xie, Lihua Wang, Andrew L. Lemire, Arthur D. Lander, Howard Y. Chang, Zhe J. Liu

## Abstract

Deconstructing the mechanism by which the 3D genome encodes genetic information to generate diverse cell types during animal development is a major challenge in biology. The contrast between the elimination of chromatin loops and domains upon Cohesin loss and the lack of downstream gene expression changes at the cell population level instigates intense debates regarding the structure-function relationship between genome organization and gene regulation. Here, by analyzing single cells after acute Cohesin removal with sequencing and spatial genome imaging techniques, we discover that, instead of dictating population-wide gene expression levels, 3D genome topology mediated by Cohesin safeguards long-range gene co-expression correlations in single cells. Notably, Cohesin loss induces gene co-activation and chromatin co-opening between active domains in *cis* up to tens of megabase apart, far beyond the typical length scale of enhancer-promoter communication. In addition, Cohesin separates Mediator protein hubs, prevents active genes in *cis* from localizing into shared hubs and blocks intersegment transfer of diverse transcriptional regulators. Together, these results support that spatial organization of the 3D genome orchestrates dynamic long-range gene and chromatin co-regulation in single living cells.

## INTRODUCTION

The mammalian genome within three-dimensional (3D) nucleus folds into high-order structures, including local chromatin loops, self-contacting topological associated domains (TADs), active or inactive compartments (Dixon et al., 2012; Lieberman-Aiden et al., 2009; Nora et al., 2012; Rao et al., 2014). Extensive genomic and imaging studies revealed that Cohesin is the primary molecular machinery driving the formation of loop domains and TADs (Bintu et al., 2018; Rao et al., 2017b; Schwarzer et al., 2017). The emerging picture is that the Cohesin ring extrudes the chromatin fiber and generates high probability contacts along its path until the extrusion is blocked by convergent CTCF sites at the domain boundary (Alipour and Marko, 2012; Fudenberg et al., 2016; Sanborn et al., 2015). Despite the well characterized role of Cohesin in genome organization, one converging and perplexing result is the lack of significant gene expression changes at the cell population level after Cohesin loss (Rao et al., 2017b; Schwarzer et al., 2017). Thus, the relationship between genome organization and transcriptional regulation remains under intense debates (Hsieh et al., 2020; Jiang et al., 2020; Su et al., 2020; Xie et al., 2022). On the other hand, it is well established that transcriptional activation is orchestrated by DNA binding transcription factors (TFs) and co-factors at enhancers and core promoters (Levine et al., 2014). Recent studies have shown that a large fraction of transcriptional regulators (i.e. EWI/FLI1, BRD4, YAP and Mediator) contain intrinsically disordered low-complexity domains (LCDs) which enable them to form concentrated protein hubs at genomic loci (Cai et al., 2019; Cho et al., 2018; Chong et al., 2018; Sabari et al., 2018). These protein hubs further coordinate transcription apparatus and engage RNA polymerase II (Pol II) for gene activation (Boija et al., 2018; Cai et al., 2019; Cho et al., 2018; Chong et al., 2018).

During animal development, cells undergo well-choreographed lineage segregation and morphogenesis events, which requires precise spatial and temporal control of multiple genes in single cells across the embryo (He et al., 2020; Mittnenzweig et al., 2021). Due to the limitation of traditional cell-population assays, the molecular basis and mechanisms controlling gene co-expression in single cells remain largely unexplored. Rapid advancements in single-cell sequencing and super-resolution genome imaging techniques enable the investigation of the structure-function relationship between genome organization, protein hub formation and gene regulation in single cells with high spatiotemporal resolution (reviewed in (Xie and Liu, 2021)). Here, by single cell sequencing and spatial genome imaging assays, we first discovered that Cohesin loss induces long-distance gene co-activation and chromatin co-opening between active domains in *cis* up to tens of megabase apart. Cohesin loss promotes spatial clustering of actively transcribed genes in *cis* accompanied by extensive Mediator hub fusion globally. In addition, Cohesin confines intersegment transfer of diverse enhancer and promoter binding transcriptional regulators. Together, these results suggest that Cohesin participates in a distinct layer of gene regulation that operates at much larger length scales than enhancer-promoter communication. And, instead of dictating average gene expression levels in the cell population, the main function of this layer of regulation is to safeguard finely balanced gene co-expression programs in single cells.

## RESULTS

### Cohesin prevents long-range, cross-domain gene co-activation in single cells

The strong impact of Cohesin loss on genome organization and the lack of gene expression changes when subsequently examined at the cell population level are perplexing results that instigated intense debates regarding the relationship between genome organization and transcriptional regulation (Hsieh et al., 2020; Jiang et al., 2020; Rao et al., 2017a; Schwarzer et al., 2017; Su et al., 2020; Xie et al., 2022). Our starting premise is that, because Cohesin is essential for mitosis and cell viability (Peters et al., 2008), the long-term consequence of Cohesin loss on gene regulation might be difficult, if not impossible, to assess. The immediate impact of Cohesin loss in single cells might be masked by population averaging. Specifically, a simple example is provided here to illustrate that two genomic features can display distinct statistical relationships (positive, negative or no correlation) at the single-cell level even when their average levels stay the same in the population (Figure 1A). To test this possibility, we took advantage of Smart-Single Cell RNA Barcoding (SCRB), a highly sensitive single-cell transcriptome assay (Bagnoli et al., 2018; Cembrowski et al., 2018; Wang et al., 2021b) to evaluate gene expression changes upon Cohesin removal (Figure 1B). Using a well-characterized mouse embryonic stem (ES) cell line with Cohesin subunit RAD21 fused to the auxin-induced degron (AID) system (Nishimura et al., 2009; Xie et al., 2022), we generated Smart-SCRB data under control and acute Cohesin loss conditions for two biological replicates. We were able to detect read fragments for ∼6,191 genes in single cells and ∼26,169 genes across populations (Figure S1A). Consistent with previous reports (Rao et al., 2017a; Schwarzer et al., 2017), we found that aggregated sequencing data at the cell population level only show subtle gene expression changes (Figure 1C and S1B).

**Figure 1.**
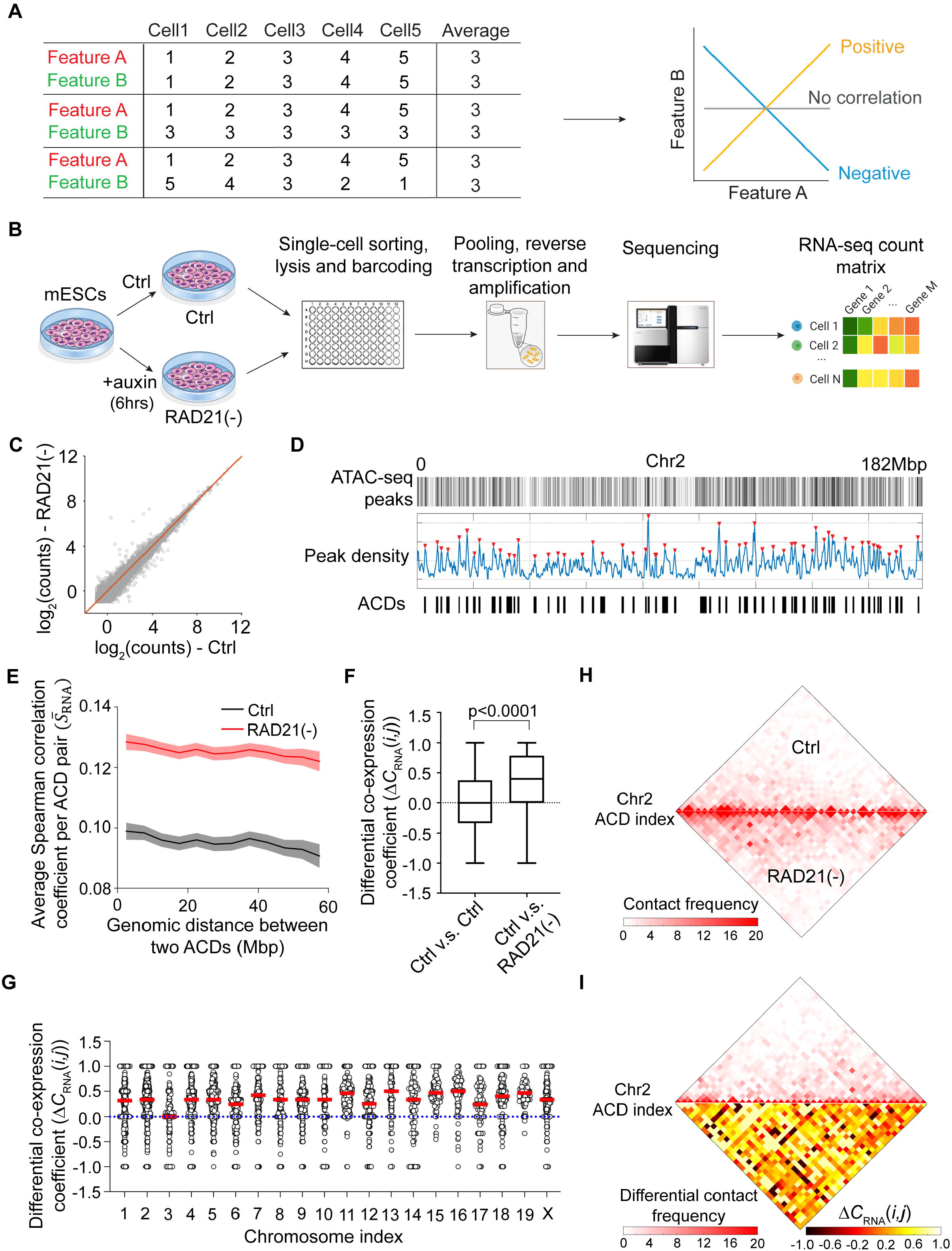
Cohesin prevents long-range, cross-domain gene co-expression in single cells. (A) A schematic diagram showing that the correlation of two genomic feature A and B at the single cell level may vary drastically even when their average levels remain the same in the cell population. (B) The workflow for Smart-SCRB sequencing of control and RAD21-depleted mESCs. (C) Acute Cohesin depletion only results in subtle changes in average gene expression at the cell population level. Each circle represents a detected gene. Average gene expression was quantified by calculating the logarithmic value of averaged sequencing counts of ∼400 individual cells. (D) Accessible chromatin domains (ACDs) were identified based on ATAC peak densities by gaussian peak fitting and thresholding for each chromosome. A total number of 776 ACDs were identified across the genome. (E) Acute Cohesin removal increases the average Spearman correlation coefficient 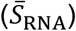 per ACD pair globally. 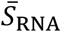 under control and Cohesin-depletion conditions was plotted as a function of the genomic distance between ACD pair after a five-point smoothing. Only the data from ACD pairs within the same chromosome were used to generate the plot. Shadow regions indicate standard error (S.E.) of the curves. (F) Box plots show the pooled statistics of differential co-expression coefficient (*ΔC*_RNA_*(i,j)*) per ACD pair throughout the whole genome after Cohesin depletion from two independent experiments. Differential co-expression coefficient calculated from two repeats with control ES cells was included as a control. The measurement was repeated twice and the non-parametric Wilcoxon test was used for statistical testing. The upper and lower whiskers represent maximum and minimum values; the box represents the range from 25% to 75% percentile; the center line represents the median; the dotted line indicates the zero-change line. (G) The statistics of *ΔC*_RNA_*(i,j)* per ACD pair before and after Cohesin depletion in chromosome 1-19 and X. Each circle indicates the value of differential co-expression coefficient for one ACD pair. The red line indicates the median value; the dotted line indicates the zero-change line. (H) Cohesin loss (lower) increases normalized contact frequency per ACD pair compared with the control condition (upper) in Chr 2. The normalized contact frequency per ACD pair was calculated by averaging the Hi-C contact frequencies over the regions within the boundaries of two ACDs. (I) Heatmaps show elevated differential co-expression coefficients per ACD pair (lower) and increased normalized differential Hi-C contact frequencies (upper) in Chr2 after Cohesin depletion. Normalized differential contact frequency per ACD pair was calculated by subtracting the normalized contact frequency in control condition from that in Cohesin-depleted condition.

Genomic studies showed that Cohesin loss tilts the balance of genome organization from TAD formation to compartmentalization, promoting cooperative multi-way chromatin interactions within active chromatin (Rao et al., 2017a; Schwarzer et al., 2017). Consistent with these findings, super-resolution imaging of the accessible genome by 3D ATAC-PALM revealed that ATAC-rich regions in the genome are organized into non-uniformly distributed clusters or accessible chromatin domains (ACDs) in the nucleus (Xie et al., 2020). Cohesin loss leads to extensive spatial mixing of ACDs (Xie et al., 2022). These results prompted us to examine whether Cohesin loss would affect co-expression correlation of genes residing in distinct ACD pairs across the genome. To evaluate cross-ACD gene co-expression, we first identified ACDs based on the local enrichment of ATAC-seq peaks in mouse ES cells. As a result, each chromosome is segmented into a string of ACDs interspaced by ATAC-poor regions (Figure 1D). In total, 776 ACDs with a size distribution from ∼0.5 to ∼2.5 Mbps were identified across the mouse genome (Figure S1C; Table S1). By analyzing gene-to-gene correlation at the single cell level using a modified, corrected Pearson residual model, we found that correlated gene expression much more likely occurs within ACDs compared with random contiguous blocks with similar gene numbers in the genome. The effect is greater for positive correlations than for negative ones (Figure S1D-E). These results are consistent with the fact that ACDs are enriched with *cis*-regulatory elements, particularly regions that positively regulate transcription (*i*.*e*. enhancers and promoters).

Then, we devised a statistical model based on Spearman’s rank-order correlation to quantify gene co-expression per ACD pair across the genome (Figure S1F-G). We found that the average co-expression Spearman correlation 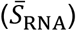 per ACD pair decays as a function of the genomic distance and Cohesin loss increases the correlation globally across all genomic length scales that we examined (up to tens of megabase) (Figure 1E). In addition, although Cohesin loss alters differential co-expression coefficient (*ΔC*_RNA_*(i,j)*) per ACD pair both positively and negatively for all chromosomes that we tested, the *ΔC*_RNA_*(i,j)* averages show uniform upward shifts toward the positive direction (Figure 1F and 1G). As a control, we performed cross-validation by using two independent scRNA-seq datasets under unperturbed conditions and revealed no significant changes (Figure 1F). Next, we computationally parsed Smart-SCRB data based on cell cycle genes and found that acute Cohesin removal led to a slight reduction of G1 (3.8%) and S (3.7%) phase cells and an increase in G2/M (7.5%) cells (Figure S1H). It is important to note that Cohesin loss significantly increases cross-ACD gene co-expression in all cell cycle phases (Figure S1I), ruling out cell-cycle effect. Together, these findings suggest that Cohesin loss selectively increases cross-ACD gene co-expression in single cells without drastically altering global gene expression at the cell population level.

To probe the link between Hi-C data with Smart-SCRB data, we next parsed Hi-C interaction frequencies before and after Cohesin loss into 776 ACDs. As expected, Cohesin loss increases Hi-C contact frequencies between ACD pairs in *cis* (Figure 1H, 1I and S1J), reflecting enhanced compartmentation of active chromatin as previously reported (Rao et al., 2017a; Schwarzer et al., 2017). However, we did not detect significant correlation between Hi-C contact frequency and the average co-expression Spearman correlation 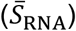 across ACD pairs (Figure S1K), probably due to the fact that Smart-SCRB only detects accumulative gene expression changes by measuring steady-state mRNA levels in single cells.

### Cohesin loss increases cross-domain chromatin co-accessibility in single cells

Next, we reason that quantifying chromatin accessibility by single-cell (sc) ATAC-seq (Buenrostro et al., 2015; Satpathy et al., 2019) would reveal immediate early regulatory changes induced by Cohesin removal. We obtained scATAC-seq data for two biological replicates before and after acute Cohesin depletion (Figure 2A). We binned scATAC-seq counts into 776 ACDs across the genome. Consistent with previous bulk ATAC-seq data showing that acute Cohesin loss causes no significant changes in chromatin accessibility at enhancers, promoters and CTCF sites at the cell population level (Xie et al., 2022), aggregated scATAC-seq data revealed little changes of chromatin accessibility in ACDs before and after Cohesin loss (Figure 2B). Then, we calculated pairwise cross-ACD chromatin co-accessibility in single cells by using a Spearman correlation model (Figure S2A). In agreement with Smart-SCRB data, we found that cross-ACD chromatin co-accessibility decays as a function of genomic distance (Figure 2C) and Cohesin loss markedly increases cross-ACD co-accessibility across all genomic length scales and all chromosomes that we examined (Figure 2C-G). As an internal reference, Cohesin loss affects little the average co-accessibility correlation between ACD pairs in *trans* (Figure S2B-C). In contrast to Smart-SCRB data, we did observe a modest but positive correlation between Hi-C contact frequency and cross-ACD co-accessibility per ACD pair (Figure S2D). The correlation becomes much stronger upon Cohesin loss (Figure S2D), implying a tighter coupling between chromatin contact and chromatin co-accessibility in the absence of Cohesin. Together, these results support a role of Cohesin in preventing cross-ACD chromatin co-opening in *cis* at the single cell level.

**Figure 2.**
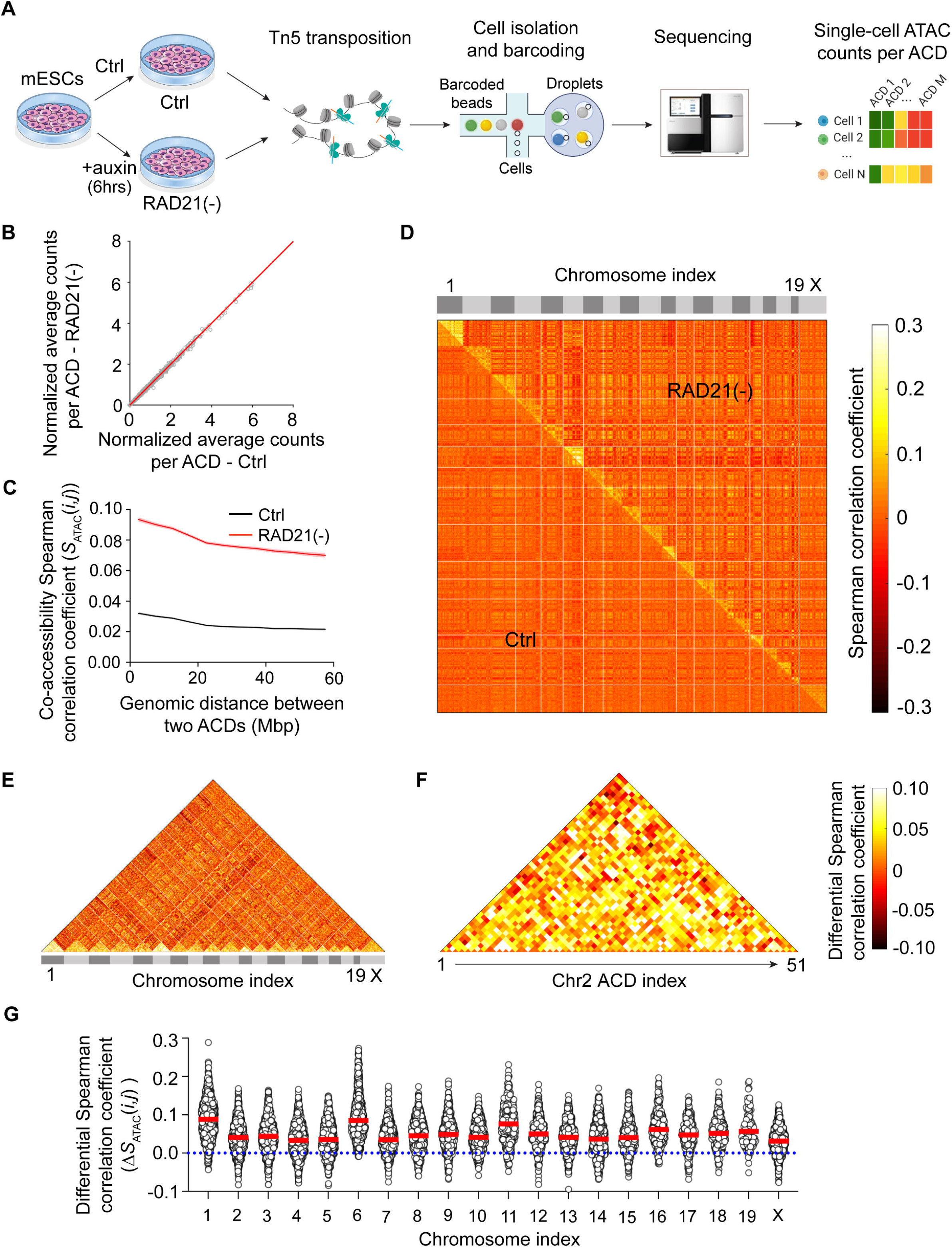
Cohesin loss increases cross-domain chromatin co-accessibility in single cells. (A) A schematic workflow for single-cell ATAC sequencing (10×Genomics) of control and RAD21-depleted mESCs. (B) Acute Cohesin loss results in no significant alteration of normalized average counts of ATAC peaks per ACD at the cell population levle. Each circle represents one ACD. Normalized average counts of ATAC peaks per ACD was calculated by dividing the accumulated ATAC-peak count per ACD by *M*_Average_ (mean ATAC-peak counts across all ACDs and cells). (C) Acute Cohesin loss increases chromatin co-accessibility per ACD pair globally. Co-accessibility Spearman correlation coefficient per ACD pair (*S*_ATAC_(*i,j*)) under control (black) and Cohesin-depleted (red) conditions was plotted as a function of the genomic distance after a five-point smoothing. Only the data from ACD pairs within the same chromosome were used to generate the plot. Shadow regions indicate standard error (S.E.) of the curves. (D) The heatmap shows that acute Cohesin loss selectively increases intra-chromosome chromatin co-accessibility per ACD pair across 20 chromosomes in the mouse genome. The calculation of cross-ACD co-accessibility Spearman correlation coefficient is described in Figure S2A. (E) The heatmap shows elevated differential co-accessibility per ACD pair within each chromosome. Differential co-accessibility Spearman correlation coefficient per ACD pair was calculated by subtracting the co-accessibility Spearman correlation coefficient value in control condition (**D**, lower left) from that in Cohesin-depleted condition (**D**, upper right). (F) The heatmap shows elevated differential co-accessibility per ACD pair in Chr2. (G) Dot plots of differential co-accessibility Spearman correlation coefficients (*ΔS*_ATAC_(*i,j*)) before and after Cohesin depletion for chromosome 1-19 and X. Every circle indicates the value of differential co-accessibility Spearman correlation coefficient per ACD pair. The red line indicates the median value, and the dotted line indicates the zero-change line.

### Cohesin prevents distant active genes in *cis* from clustering and co-activation

It was recently shown that early changes in gene activities and 3D genome organization can be investigated by genome-wide intron fluorescence in *situ* hybridization (FISH) (Shah et al., 2018). To complement with our genomic results and probe gene co-activation over long distances, we sought to use intron RNA-FISH to examine spatial distribution of actively transcribed genes across Chromosome 2 (Chr 2). Specifically, we designed single-molecule RNA-FISH probes to simultaneously target intron regions of 208 active genes across Chr 2 (Figure 3A, Table S2). 3D Airyscan imaging showed that actively transcribed genes are organized to individual, well separated puncta before Cohesin depletion (Figure 3B). Cohesin loss induced extensive clustering of these puncta, leading to the formation of larger and more connected structures in the nucleus (Figure 3B-D, Movie S1). Consistent with these findings, super resolution imaging by 3D ATAC-PALM and Oligo-PAINT DNA-FISH demonstrated that spatial mixing of accessible chromatin upon Cohesin loss mainly occurs in *cis* with accessible sites within the same chromosome moving closer to each other without substantially affecting the distance between accessible sites in *trans* (Xie et al., 2022). Together, these imaging data suggest that Cohesin supplies a broad function to prevent active genes in *cis* from clustering.

**Figure 3.**
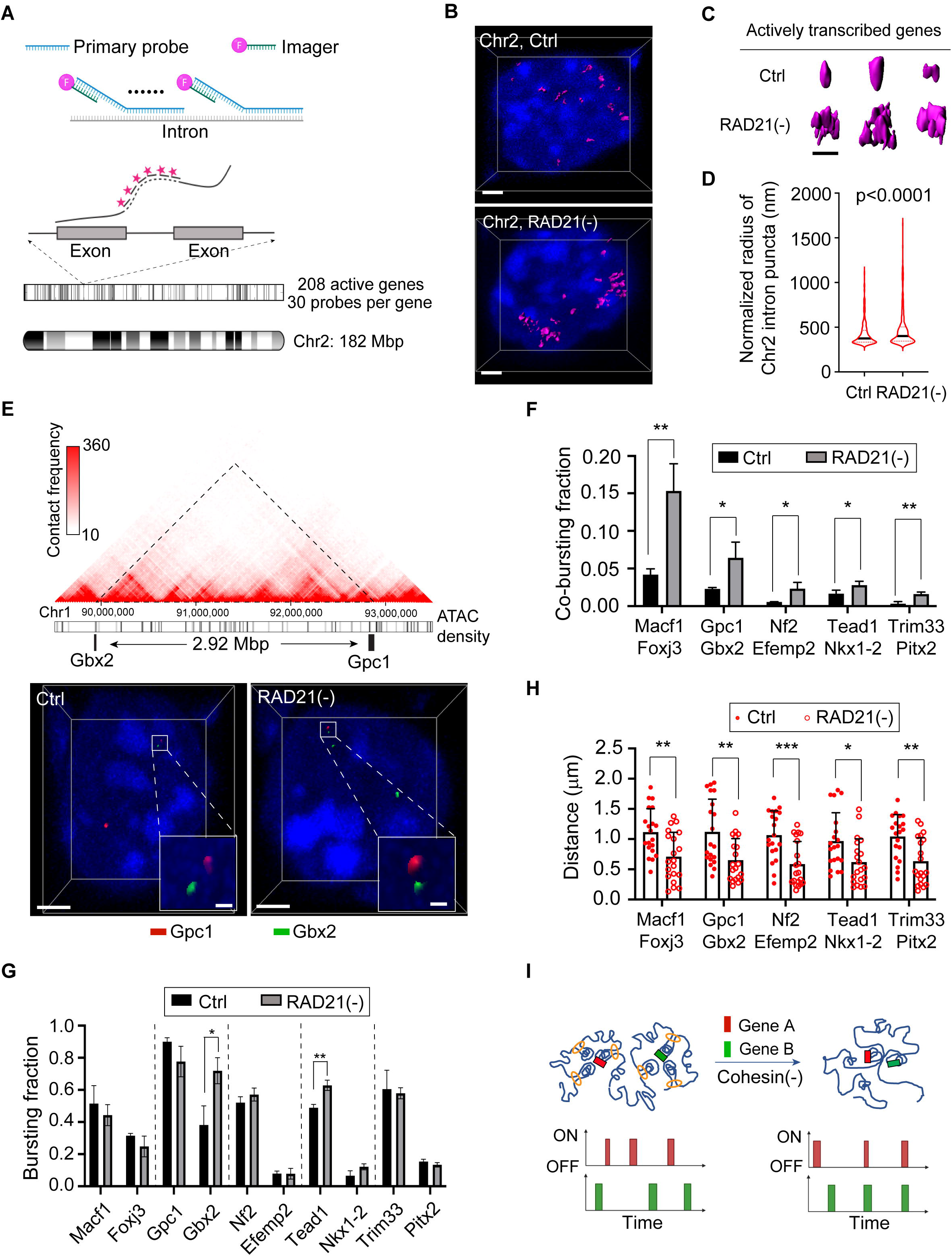
Cohesin blocks clustering and co-activation of distant active genes in *cis*. (A) Transcription bursting sites of 208 active genes from Chr2 are collectively imaged by using intron primary probes (30 probes per gene) and a global imager probe conjugated with Alexa-647N. (B) Representative 3D *iso*-surface images show transcriptional bursting sites from 208 active genes across Chr2 before (left) and after (right) Cohesin loss. Nuclei were counter-stained with DAPI (Blue). Scale bar, 2μm. (C) Isolated representative 3D *iso*-surfaces for typical intron puncta formed by transcription bursting sites of 208 genes across Chr2 before and after Cohesin loss. Scale bar, 1μm. (D) Violin plots show the normalized radii of detected Chr2 intron puncta before and after acute Cohesin depletion. Black lines are the median values and dotted lines are 25% and 75% percent quantiles. The measurement was repeated for three times and non-parametric Wilcoxon test was used for statistical testing. (E) Cohesin loss decreases the physical distance between co-activated lineage-specific genes Gbx2 (ectoderm) and Gpc1 (mesendoderm). Upper panel, genomic positions of Gbx2 and Gpc1 in Chr1 with Hi-C and ATAC density information. Lower panel, representative 3D *iso*-surface images of Gbx2 and Gpc1 bursting sites before and after Cohesin loss. Scale bar, 2μm. Inlet scale bar, 200nm. (F) Cohesin loss elevates co-bursting fractions of 5 pairs of lineage-specific genes. Error bars indicate standard deviation (S.D.). The measurement was repeated three times and Student’s t-test was used for statistical testing. *, p < 0.05; **, p < 0.01; ***, p < 0.001. (G) Cohesin loss decreases average physical distances between co-activated lineage-specific genes in *cis* (5 pairs). Error bars indicate standard deviation (S.D.). The experiment was repeated for three times and Wilcoxon test was used for statistical testing. (H) Cohesin loss does not cause uniformed up- or down-regulation of bursting fractions for individual genes. Error bars indicate standard deviation (S.D.). The measurement was repeated three times and Student’s t-test was used for statistical testing. (I) A schematic diagram showing chromatin reorganization (upper) and gene co-activation (lower) upon Cohesin depletion.

To test the potential impact of Cohesin loss on genes involved in cell fate acquisition, we next performed intron RNA-FISH to measure co-bursting fractions of 5 representative pairs of lineage specific genes located in distinct accessible regions far apart in *cis* (from 2.9 ∼ 25.9 Mbps away from each other) (Figure 3E and S3A-B). Single-cell RNA-FISH analysis showed that Cohesin loss increases the co-bursting fraction for each of the 5 gene pairs tested (Figure 3F) while bursting fractions for individual genes do not display a consistent trend of up- or down-regulation in the population (Figure 3G). These results suggest that Cohesin loss selectively increases co-activation of these gene pairs in single cells rather than alters activation frequencies of individual genes in the population (Figure 3I). We next performed gene expression simulation and showed that the increase in co-expression correlation does not perturb average gene expression levels in the population (Figure S4A-G). We also observed a reduction of average physical distances between these co-activated gene pairs (Figure 3H and S3C), echoing the finding that Cohesin loss triggers prominent clustering of actively transcribed genes in *cis* across Chr 2.

### Cohesin blocks Mediator hub fusion in live cells

It is well established that Cohesin loss leads to enhanced compartmentalization of active compartments (Rao et al., 2017a; Schwarzer et al., 2017) and spatial mixing of ACDs (Xie et al., 2022; Xie et al., 2020). However, it remains elusive whether and how these changes in genome organization affect spatial distribution of transcriptional regulators in the nucleus, particularly their ability to form protein hubs. To address this question, we decided to image and analyze Mediator hubs because it has been well established that Mediator is structurally associated with RNA Polymerase II (Abdella et al., 2021) and is essential for coordinating transcription activation genome-wide (El Khattabi et al., 2019; Kagey et al., 2010). In addition, Mediator forms protein hubs that are spatially correlated with transcriptional activities in living cells (Cho et al., 2018; Sabari et al., 2018). We utilized CRISPR/Cas9-based genome editing and fused HaloTag to endogenous Mediator subunits MED1/MED6 in the ES cell line harboring the RAD21 degron. It is worth noting that both MED1 and MED6 contain low-complexity domains (LCDs) (Figure S5A-B). The bi-allelic fusion of MED1 or MED6 with HaloTag did not significantly affect cell cycle and viability of ES cells (Figure S5C-I). Consistent with previous reports (Cho et al., 2018; Sabari et al., 2018), we confirmed that MED1/MED6-HaloTag form concentrated hubs in both fixed and live cells (Figure 4A and 4D, Movie S2 and S3). Multiplex RNA-FISH and protein imaging also showed that actively transcribed gene is co-localized with Mediator hubs but is spatially separate from heterochromatic regions labeled by HP1-GFP (Figure S6A-B).

**Figure 4.**
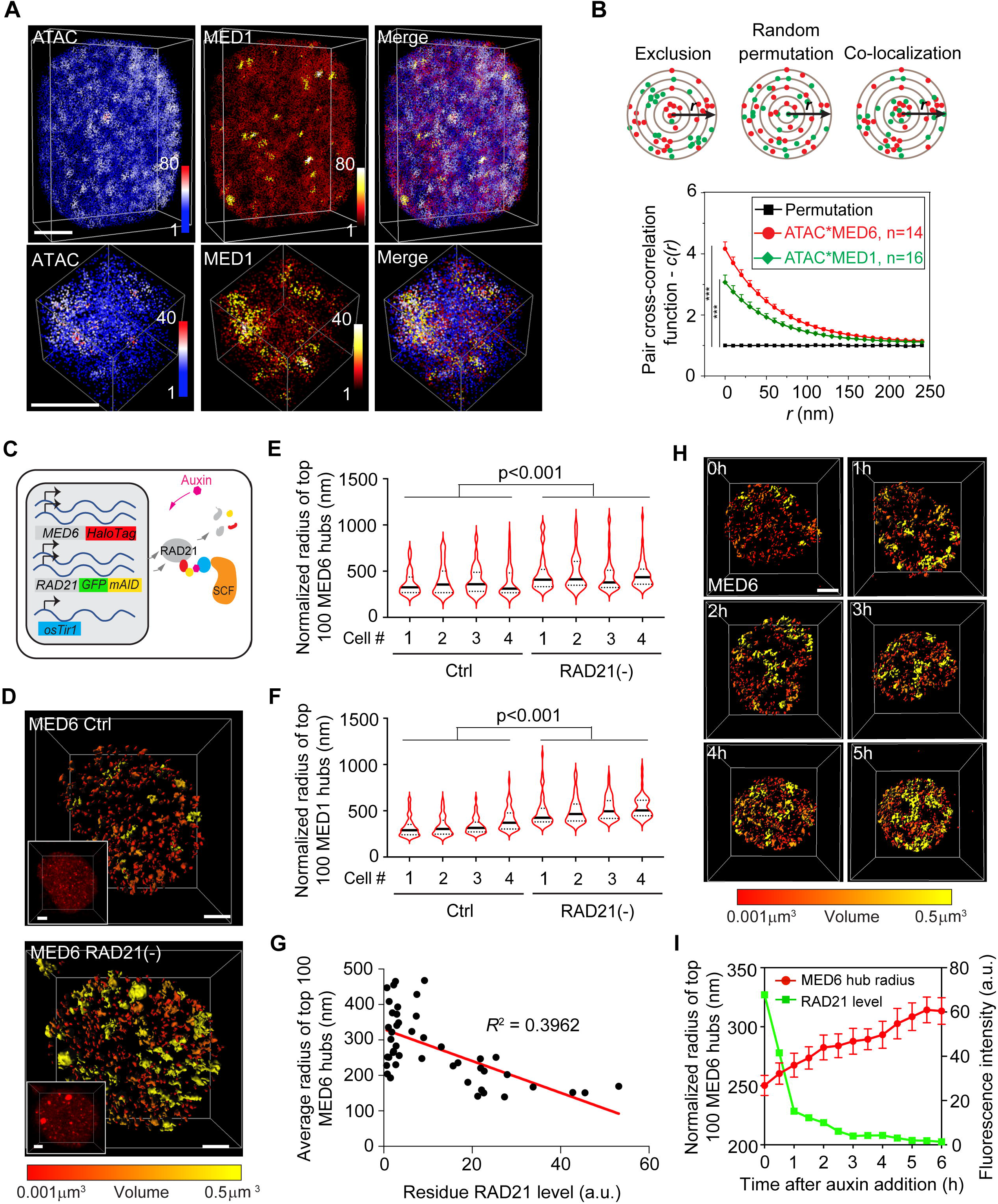
Cohesin maintains physical separation between Mediator hubs. (A) Two-color 3D PALM imaging captures spatial distribution of both accessible chromatin sites (ATAC) (left) and MED1-HaloTag (middle) localizations. Color bars indicate localization densities. Scale bar, 2μm. See Movie S2 for 3D rotatory rendering. In the lower panel, the cropped localization map indicates that ACDs colocalize with MED1 hubs. Color bars indicate localization densities. Scale bar, 500nm. (B) Quantification of colocalization of accessible chromatin localizations and MED1 (or MED6) localizations by pair cross-correlation function *c*(*r*). Error bar indicates the standard error (S.E.) of measured values. In the upper panel, schematic shows three different spatial relationship – exclusion, uncorrelated (random permutation) and colocalization – between two localization maps. The experiment was repeated for three times and non-parametric Wilcoxon test was used for statistical testing. (C) The diagram shows biallelic integration of HaloTag and GFP-mAID into endogenous *Med6* and *Rad21* gene loci. Auxin triggers rapid degradation of RAD21 through ubiquitination and proteosome mediated degradation. (D) 3D *iso*-surface reconstruction of MED6 hubs before (upper) and after (lower) Cohesin depletion. Figure inlets show Med6 fluorescence images used for the 3D reconstruction (rendered by Imaris). The Mediator hubs are color-coded based on their 3D volumes as indicated by the color bar below. Scale bar, 2μm. Inlet scale bar, 2μm. **(E-F)** Violin plots show the size distribution of MED6 (**C**) and MED1 (**D**) hubs before and after Cohesin depletion in single cells. Black lines are the median values and dotted lines are 25% and 75% percent quantiles. Non-parametric Wilcoxon test was used for statistical testing. (G) MED6 hub sizes are inversely correlated with residual RAD21 protein levels in single cells. (H) Time-lapse imaging of MED6 hubs during acute Cohesion depletion. Images were taken every 30 mins for 6 hrs upon the auxin treatment. MED6 hubs are color-coded based on their 3D volumes as indicated by the colormap below. Scale bar, 2μm. (I) Quantification of MED6 hub sizes (red) and RAD21 residual levels (green) from time-lapse imaging volumes. Error bar indicates the standard error (S.E.) of normalized radius of top 100 MED6 hubs.

To investigate the spatial relationship between ACDs and Mediator hubs, we next performed multiplexed 3D ATAC and Mediator PALM imaging. Super-resolved images showed that Mediator hubs co-localize with ACDs across the nucleus (Figure 4A and S6C, Movie S2). 3D visualization and pair cross-correlation analysis of individual regions with co-existing clusters indicated strong spatial correlation between signals from these two channels (Figure 4B and S6D), suggesting that Mediator hubs overlap with ACDs extensively. It is worth noting that, in stark contrast to the previous result that the formation of ACDs is insensitive to α-Amanitin treatment and thus is independent of transcription (Xie et al., 2022), α-Amanitin treatment drastically reduced the number of Mediator hubs in both control and Cohesin depleted cells (Figure S6E-G), suggesting that Mediator hub formation is dependent on transcription and is likely downstream of ACD formation.

Next, we coupled acute depletion of Cohesin with high-resolution 3D imaging of Mediator hubs in live cells by Airyscan microscopy (Figure 4C-D). We found that acute Cohesin removal induces extensive structural changes of Mediator hubs with a significant increase of the average hub size (Figure 4D-F; Movie S3). Importantly, the average size of Mediator hubs is inversely correlated with residual RAD21 levels in single cells (Figure 4G), suggesting a dose-dependent effect. There is also an increase in the formation of “super hubs” (*r* > 500 nm) upon Cohesin removal by dynamic fusion of smaller hubs observed under live imaging (Figure 4H-I; Movie S4). It is worth noting that acute Cohesin depletion does not affect the nucleus size (Xie et al., 2022) nor MED1/MED6 protein levels (Figure S5E-F) excluding the possibility that mediator hub fusion is induced by altered protein concentrations. Together, these results suggest that Cohesin maintains physical separation between Mediator hubs.

### Mediator hub fusion spatially correlates with aberrant gene co-expression

Recent studies have shown that protein hubs formed by transcription factors and cofactors are correlated with transcriptional activities (Cai et al., 2019; Cho et al., 2018; Chong et al., 2018; Sabari et al., 2018). To investigate the spatial relationship between Mediator hubs and gene co-expression, we first analyzed the 3D volume overlap between Mediator hubs and intron-FISH puncta formed by actively transcribed genes in Chr 2. We found that Mediator hub fusion spatially correlates with clustering of intron puncta across Chr 2 upon Cohesin loss (Figure 5A-C), suggesting that more active genes are localized into larger, more connected mediator hubs. Next, we performed Mediator hub imaging and intron-RNA-FISH of the 5 pairs of lineage specific genes that we analyzed previously (Figure 3E and S3A). We quantified the fraction of co-activated genes localized to shared Mediator hubs (Figure 5D) and found significantly higher co-localization frequencies upon Cohesin loss (Figure 5E). As a control, we analyzed co-localization between Mediator hubs and co-activated gene pairs localized within the same ACDs and found no significant increase in the colocalization frequencies (Figure 5E). Together, these results showed that Cohesin not only spatially separates distant active genes in *cis* but also prevents them from localizing to shared Mediator hubs.

**Figure 5.**
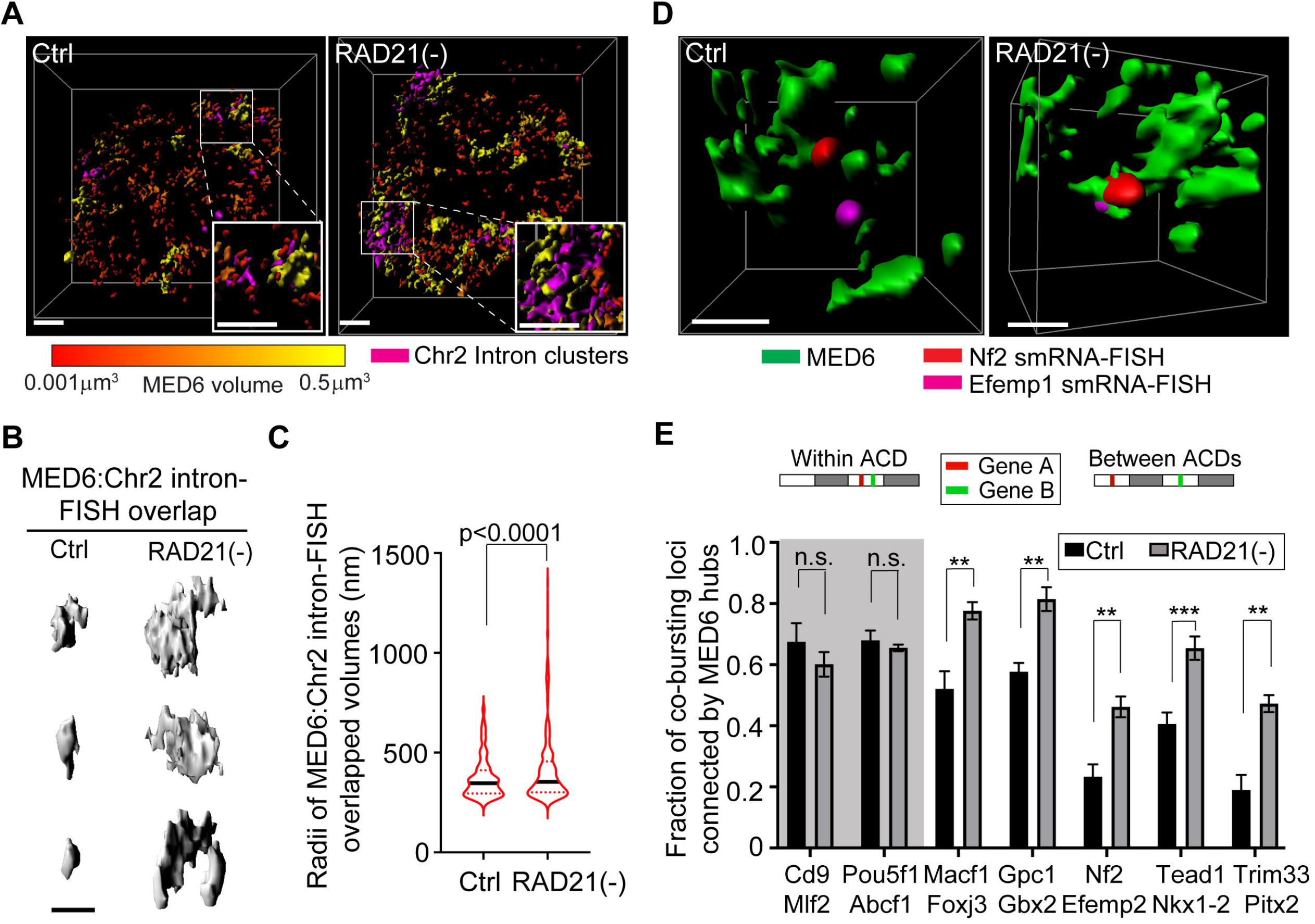
Cohesin prevents actively transcribed genes in *cis* from localizing to shared Mediator hubs. (A) 3D *iso*-surface rendering of overlaps between MED6 hubs (volume is color-coded) and transcription bursting sites for 208 genes in Chr2 (magenta) before and after RAD21 depletion. MED6 hubs are color-coded based on their 3D volumes as indicated by the colormap below. Scale bar, 2μm. Inlet scale bar, 2μm. (B) Representative 3D volume overlaps between MED6 hubs and Chr2 intron clusters before and after Cohesin loss. Scale bar, 1μm. (C) Violin plots show the statistics of 3D volume overlaps between MED6 hubs and transcription bursting sites for 208 genes in Chr2 before and after Cohesin loss. Black lines are the median values and dotted lines are 25% and 75% percent quantiles. The measurements were obtained from 20 cells and non-parametric Wilcoxon test was used for statistical testing. (D) 3D *iso*-surface reconstruction of MED6 hubs and intron-FISH signals showing a representative case that co-bursting gene loci are connected by MED6 hubs after Cohesin loss. Scale bar, 500nm. (E) Bar plots and statistics for the fraction of co-bursting loci connected by MED6 hubs for two pairs of genes within the same ACD (shadowed) and five gene pairs cross ACDs before and after Cohesin loss. The fraction was calculated by dividing the number of co-bursting loci which share a common MED6 hub by the total number of co-bursting loci. The measurement was repeated three times and Student’s t-test was used for statistical testing. *, p < 0.05; **, p < 0.01; ***, p < 0.001.

### Cohesin confines intersegment transfer of diverse transcriptional regulators

The increase in cross-ACD chromatin co-opening, the reduced distance between co-activated genes and their co-localization into shared mediator hubs further prompt us to investigate whether these functional and structural changes upon Cohesin loss are associated with altered transcription factor binding and diffusion dynamics in live cells. To address this question, we first selected a broad range of transcriptional regulators representing distinct processes in transcriptional activation. These factors include site-specific transcription factors (OCT4), histone mark reader (BRD4), Mediator subunits (MED1 and MED6) and core promoter factor (TATA-binding protein (TBP). Histone H2B and H2A.Z were also included as general markers for chromatin and active chromatin, respectively. We used bright, cell-permeable Janelia fluorophore dye-JF549 (Grimm et al., 2015) to label HaloTag fusion proteins that were either expressed endogenously (MED1, MED6 and BRD4) or were stably expressed at relatively low levels (OCT4, TBP, H2B and H2A.Z) in ES cells (Figure S5E-F and S7A-B). We first performed fast live-cell, single-molecule imaging (100Hz) followed by automated single molecule tracking and diffusion analysis as previously described (Figure S7C) (Chen et al., 2014b; Liu et al., 2018a; Liu et al., 2014; Piccolo et al., 2019). Surprisingly, a simple two- or three-state model (Hansen et al., 2018) that assumes a bound state and one or two diffusive states failed to reveal significant differences in histone and TF dynamics before and after Cohesin loss (Figure 6A-C and S7D-G), suggesting that Cohesin loss does not notably affect their apparent diffusion and binding equilibria in the nucleus.

**Figure 6.**
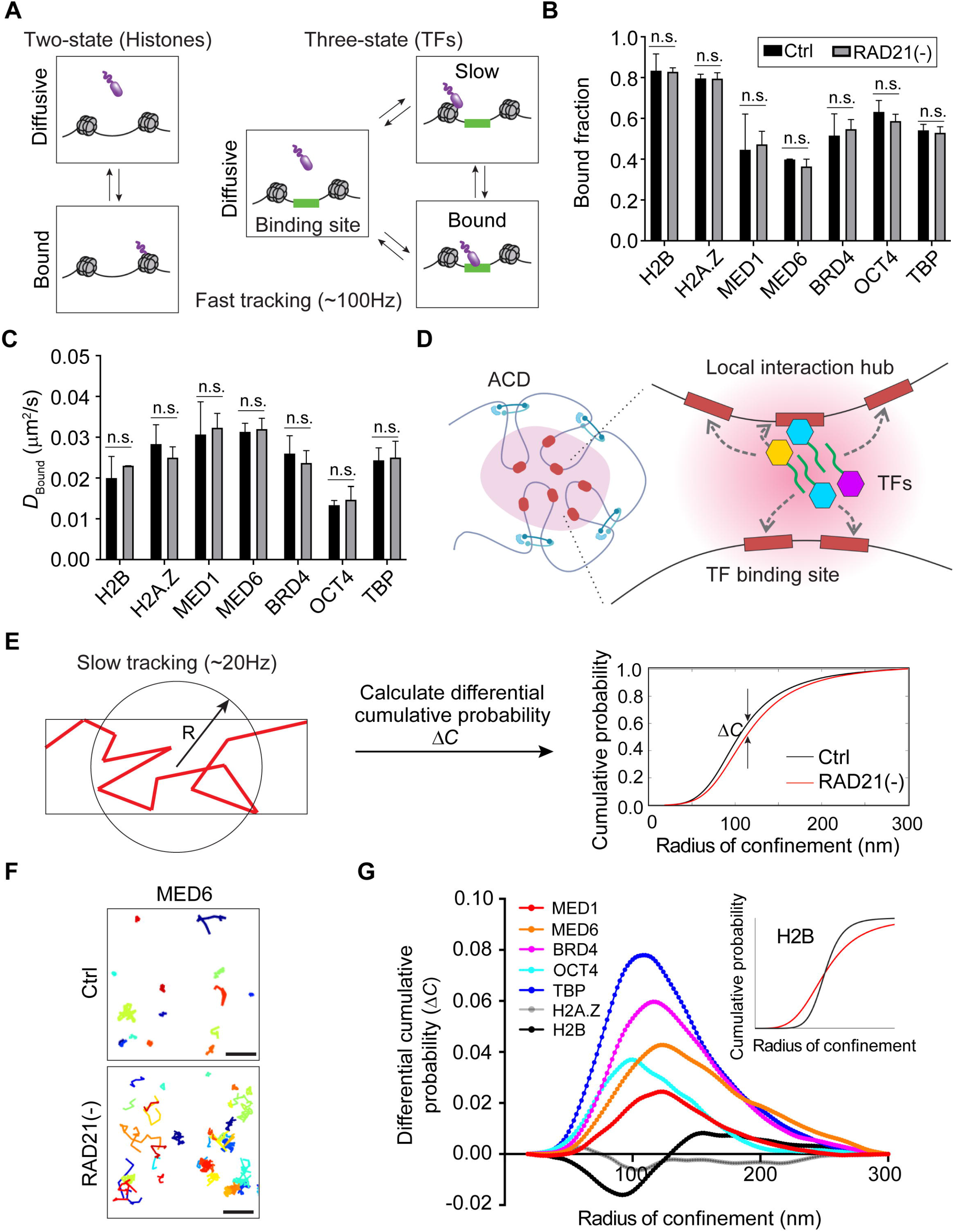
Cohesin confines intersegment transfer of diverse transcriptional regulators. (A) The diagram shows two-state and three-state fitting of single-particle tracks for histone subunits and transcriptional regulators, respectively. For this analysis, fast single molecule imaging (100Hz) was used to capture the movement of both diffusive and bound molecules. (B) Fast tracking reveals that Cohesin loss does not significantly alter apparent bound fractions for histone subunits (H2B and H2A.Z) and a broad range of transcriptional regulators (MED1, MED6, BRD4, OCT4 and TBP). Error bars indicate standard deviation (S.D.). The measurement was repeated for three times and Student’s t-test was used for statistical testing. n.s., not significant. (C) Fast tracking reveals Cohesin loss has little effect on apparent diffusion coefficients for histone subunits (H2B and H2A.Z) and a broad range of transcriptional regulators (MED1, MED6, BRD4, OCT4 and TBP) in the bound state. Error bars indicate standard deviation (S.D.). The measurement was repeated for three times and Student’s t-test was used for statistical testing. n.s., not significant. (D) Transcription factors search for targets in a local interaction hub or condensate by a bound-state dominant mode. Intersegment transfer via facilitated diffusion is proposed to be guided by transient interactions with other co-regulators (protein or noncoding RNAs) in the hub. (E)The diagram illustrates the quantification of radius of confinement (RC) and the calculation of its differential cumulative probability *ΔC* (*ΔC= ΔC*_Ctrl_*-ΔC*_RAD21(-)_) over analyzed tracks. For this analysis, both loosely and tight bound fractions were obtained by using a slower frame rate (20Hz). (F) Examples of MED6 single molecule trajectories (colored by random colors) before and after Cohesin loss obtained by slow tracking. Scale bar, 500 nm. (G) Differential cumulative probability (*ΔC*) of RC for histone subunits (H2B and H2A.Z) and a broad range of transcriptional regulators (MED1, MED6, BRD4, OCT4 and TBP). Histone subunits H2B and H2A.Z were analyzed as controls. Inlet, schematic curves showing cumulative probability of RC for H2B before (black) and after (red) Cohesin loss.

Next, we analyzed the transfer of transcriptional regulators between binding sites in the nucleus – a processed known as intersegment transfer (Figure 6D). Intersegment transfer via facilitated diffusion has been proposed as a mechanism underlying long-distance enhancer-promoter communication (Alexander et al., 2019; Karr et al., 2022; Liu et al., 2014; Liu and Tjian, 2018; Mir et al., 2018; Wollman et al., 2017). Specifically, it was shown that TF searches for targets within local interaction hubs via a bound-state dominant mode (Liu et al., 2018b; Liu et al., 2014; Mir et al., 2017) likely guided by transient interactions with cofactors and noncoding RNAs (Bhat et al., 2021; Quinodoz et al., 2021). Recently, a parameter called the Radius of Confinement (RC) was specifically developed to analyze intersegment transfer by quantifying the degree of confinement for TFs that are both tightly and loosely bound to chromatin (Lerner et al., 2020). By using a longer acquisition time (50ms) and thus selectively imaging the dynamics of chromatin-bound molecules, we compared the RC cumulative probability distributions before and after Cohesin loss (Figure 6E). We found that Cohesin loss consistently increases RC for each representative transcriptional regulator tested (Figure 6F-G). On the contrary, the RC for the active chromatin marker H2A.Z shows slight decreases (Figure 6G). Interestingly, the RC curve for the general chromatin marker H2B adopts a less sigmoid shape (Figure 6G), implying more homogeneous chromatin movements upon Cohesin loss. Together, these results suggest that Cohesin loss has distinct effects on transcriptional regulators and chromatin and that Cohesin likely plays a role in sequestering intersegment transfer of a broad range of enhancer and core promoter binding transcriptional regulators.

## DISCUSSION

### Cohesin orchestrates long-range gene co-regulation in single cells

Nucleus is a crowded environment consisting of high densities of chromatin and a myriad of chromatin binding proteins involved in diverse biological activities (transcription, replication, DNA repair, etc). Due to limitations associated with traditional cell-population end-point assays such as biochemistry and genetics, most studies in past decades focus on dissecting the mechanism underlying activation of single genes within specific cell populations. These studies were fruitful at revealing a cohort of transcription activators and co-activators that function at the enhancer and the core promoter level to modulate Pol II activities. Loss of function in this layer of regulation usually leads to global gene expression changes in specific cell populations (Levine et al., 2014; Levine and Tjian, 2003; Mannervik et al., 1999). However understandably, due to lack of resolution to dissect molecular mechanisms at the single cell level, these cell-population based assays fell short of revealing mechanistic insights into co-regulation of multiple genes in single cells.

During each cell cycle, the overall chromatin structure is altered drastically accompanied by cell-cycle dependent shifts in gene activities. Thus, the proper folding and organization of the 3D genome has been regarded as the foundation to other regulatory activities throughout the cell cycle. The role of Cohesin in genome organization has long been established. During prometaphase, Cohesin is concentrated at centromeres and is responsible for holding sister chromatids together (Peters et al., 2008). Recently, a combination of acute Cohesin depletion and Hi-C genomic assays revealed that Cohesin loss eliminates all loop domains and TADs but enhances chromatin compartmentalization in the interphase nucleus (Kim et al., 2019; Rao et al., 2017b; Schwarzer et al., 2017). Despite the seminal role of Cohesin in genome organization, one surprising result is that Cohesin loss only leads to subtle gene expression changes at the cell population level (Rao et al., 2017a; Schwarzer et al., 2017), instigating debates on the role of genome organization in gene regulation (Hsieh et al., 2020; Jiang et al., 2020; Su et al., 2020; Xie et al., 2022).

Here, we reason that the immediate impact of Cohesin loss on gene regulation might be masked by population averaging. Thus, we used single-cell sequencing techniques to analyze gene and gene regulatory activities in single cells. Consistent with previous reports, we found that cell-population ensemble of scRNA-seq data reveals little changes in global gene expression, suggesting that Cohesin does not participate in a rate limiting step of determining average gene expression levels at the cell population level. However, single-cell analysis revealed that Cohesin loss induces extensive gene co-activation and chromatin co-opening between active domains in *cis* up to tens of megabase apart, far exceeding the typical length scale of enhancer-promoter interactions (mostly in the kilobase and rarely into the megabase range) (Levine et al., 2014). These results suggest that Cohesin participates in a distinct layer of gene regulation that operates at much larger length scales than enhancer-promoter communication. And, instead of dictating average gene expression levels in defined cell populations, the main function of this layer of regulation is to safeguard finely balanced long-range gene co-expression correlations in single cells. In addition, spatial genome and protein imaging in single cells reveals that Cohesin loss leads to clustering of actively transcribed genes in *cis* accompanied by extensive mediator hub fusion and extended TF exploration range via intersegment transfer (Figure 7A-B). These results suggest that Cohesin may exert its role in gene regulation by modulating spatiotemporal distribution and dynamics of biochemical reactants (chromatin and protein) in the nucleus.

**Figure 7.**
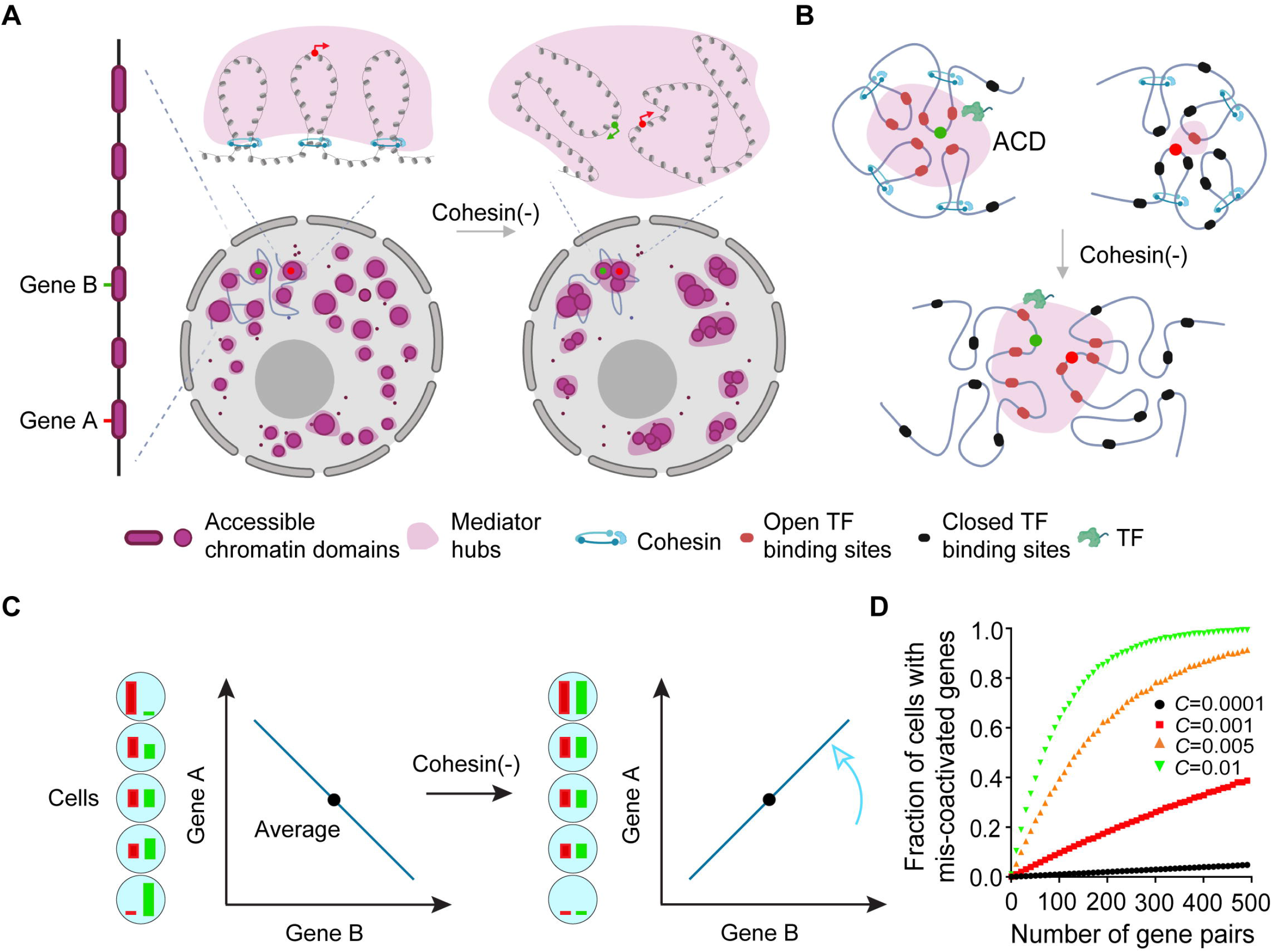
A model depicting the role of Cohesin in genome organization and gene regulation. (A) Cohesin loss leads to disruption of chromatin loops, spatial mixing of accessible chromatin domains and fusion of mediator hubs. As a result, the average distance between distant active genes (green and red dots) in *cis* decreases accompanied by colocalization of the two genes to into a shared Mediator hub and an elevated chance for their co-activation. (B) Cohesin loss leads to spatial mixing of ACDs, Mediator hub fusion and correlated cross-ACD chromatin co-opening. As a result, TF can explore large areas via intersegment transfer. (C) A schematic diagram illustrating how the increased co-expression correlation upon Cohesin loss would influence relative expression levels of gene A (red bars) and B (green bars) in single cells without affecting their average expression levels (black dots) in the population. (D) Simulation results showing the fraction of cells carrying mis-coactivated genes as a function of the number of co-activated gene pairs (*x*-axis) with a range of co-expression coefficients (*C*, color-coded).

Looking back from this point, we found that lack of gene expression changes at the cell population level upon Cohesin loss is the major unexpected result that led to the widespread speculation that 3D genome organization might be functionally decoupled from gene regulation (Hsieh et al., 2020; Jiang et al., 2020; Rao et al., 2017a; Schwarzer et al., 2017; Su et al., 2020; Xie et al., 2022). It is worth noting that ATAC-rich ACDs can be robustly detected in bulk and in single cells by genomics and imaging assays (Xie et al., 2022; Xie et al., 2020). Cohesin loss induces correlated structure-function changes across ACDs, suggesting ACDs as operational gene regulatory units in *situ*.

### Physical mechanisms by which Cohesin regulates cross-ACD gene co-expression

It is well-established that gene activation is orchestrated by transcription factors and co-factors that are dynamically assembled at the enhancer and the core promoter to modulate transcription initiation and elongation. Emerging studies indicated that a large number of transcriptional regulators contain low complexity domains which enable them to form protein hubs to further coordinate transcription apparatus and engage RNA polymerase II (Pol II) for gene activation (Boija et al., 2018; Cho et al., 2018; Sabari et al., 2018; Wang et al., 2021a). Thus, to understand the role of Cohesin in gene regulation, it is essential to examine how Cohesin regulates 1) the spatial organization of *cis*-regulatory elements (*i*.*e*. enhancers and promoters), 2) the exchange and assembly dynamics of transcriptional regulators on these elements and 3) the ability of transcription factors to form protein hubs.

The first informative clue comes from the recent report showing that acute Cohesin removal led to extensive spatial mixing and enhanced clustering of active chromatin, reducing the average distance between accessible sites and *cis*-regulatory elements globally (Xie et al., 2022). This result suggests that super enhancers (clusters of enhancers in the linear genome) previously shown to drive global gene activation (Hnisz et al., 2013; Whyte et al., 2013) would also move closer to each other upon Cohesin loss. Here, we found that the spatial mixing of active chromatin upon Cohesin loss is accompanied by clustering of active genes in *cis* and extensive fusion of Mediator hubs. As a result, genes located in distant active domains have much higher chance of localizing to shared Mediator hubs. It has been well demonstrated that Mediator is an integral part of the transcription initiation machinery and coordinates gene expression genome-wide (El Khattabi et al., 2019; Kagey et al., 2010; Wollman et al., 2017). In addition, Mediator hubs are correlated with transcriptional activities in living cells (Cho et al., 2018). Thus, these structural changes in ACDs and Mediator hubs could potentially subject genes across architectural boundaries under the influence of similar local gene regulatory fluxes, elevating the chance of their co-activation. Consistent with this model, Cohesin loss promotes cross-ACD chromatin co-accessibility in *cis*, indicating that Cohesin prevents chromatin remodeling activities from simultaneously acting on distant ACDs (Figure 7B). Indeed, live-cell, single-molecule imaging show that Cohesin inhibits intersegment transfer of diverse transcriptional regulators over long distances. Together, these results suggest that, at least, three non-exclusive and possibly interlinked mechanisms underlie the ability of Cohesin to regulate cross-domain gene co-expression, specifically by 1) physically separating ACDs, 2) blocking the subsequent fusion of protein hubs and 3) inhibiting intersegment transfer of transcriptional regulators (Figure 7A-B). It is worth noting that our results also suggest that the formation of ACDs is functionally upstream of that of Mediator hubs, because the formation of ACDs is insensitive to α-Amanitin treatment and thus is independent of transcription (Xie et al., 2022) while α-Amanitin treatment drastically reduced the number and size of Mediator hubs. Together, our results provide insights into potential transmission mechanisms by which Cohesin-mediated genome organization influences gene co-regulation in single cells.

### The connection between genome organization and cell fate determination

In a developing embryo, cells undergo well-choreographed lineage specification and morphogenesis events, which requires precise spatiotemporal control of multiple genes in single cells across the embryo. This premise underscores the necessity for higher organisms to implement mechanisms for coordinating genes genome-wide to co-express at desired relative levels in single cells. Here, our data suggest that Cohesin-mediated genome organization participates in controlling long-distance gene co-expression and chromatin co-accessibility correlations in single cells, which could in turn feedback to influence collective cellular behaviors. For instance, crucial developmental processes such as organogenesis and morphogenesis are usually driven by complex gene expression gradients across the embryo. Altering gene co-expression correlations could perturb relative arrangements between these gradients (Figure 7C) and thus disrupt finely balanced developmental programs. Interestingly, simulation shows that when propagating unregulated gene co-activation to multiple gene pairs in the genome just as shown here by scRNA-seq and intron-FISH experiments, gene co-expression defects arise rapidly in large cell fractions (Figure 7D). These results underscore the physiological necessity for precise regulation of gene co-expression at the single cell level. Most mutations associated with a human genetic disease named Cornelia *de* lange syndrome (C*d*LS) occur on proteins facilitating the loading of the Cohesin ring onto chromatin or on the Cohesin ring itself. The phenotypes of C*d*LS include cell lineage assignment errors and developmental retardation (Kline et al., 2007). In fact, analyses of Cohesin loading factor-Nipbl deficient tissues and cells from Drosophila, mouse, zebrafish and man all suggest that C*d*LS birth defects are caused by subtly mis-regulating the expression of large numbers of genes (Kawauchi et al., 2016; Muto et al., 2011; Santos et al., 2016; Weiss et al., 2021; Wu et al., 2015). One reasonable hypothesis deduced from the collective findings here is that these disease phenotypes could be at least partially due to unbalanced, stochastic alternation of gene co-expression correlations in single cells which would in turn compromise the efficiency and stability of cell fate acquisition during development. Future studies within specific developmental contexts and disease states may gain more insights into the functional link between genome organization, gene regulation and cell-fate choices.

## METHODS

### Chemicals

The plant auxin analog Indole-3-acetic acid (IAA) sodium salt (Millipore Sigma, Cat. I5148) was dissolved in ddH_2_O with a stock concentration of 500mM, aliquoted and stored at −20°C, and used at the final concentration of 500µM (acute depletion) or 5µM for 6 hrs. α-Amanitin (Tocris, Cat. 4025) was dissolved at 1mg/mL stock in ddH_2_O, aliquoted and stored at −20°C, and used at the final concentration of 100µg/ml.

### Cell culture

JM8.N4 mouse embryonic stem cells (mESCs) from the C57BL/6N strain and their genome-edited derivatives were routinely cultured in 60mm plates coated with 0.1% gelatin without feeders at 37°C and 5% CO_2_. The mESC culture medium was composed of the optimized knockout DMEM for mESCs (Thermo Fisher Scientific, Cat. 10829-018), 15% ESC-qualified Fetal Bovine Serum (ATCC, Cat. SCRR-30-2020), 1000 units of Leukemia inhibitory factor (LIF) (self-purified), 1mM GlutaMAX (Thermo Fisher Scientific Cat. 35050-061), 0.1mM MEM nonessential amino acids (Thermo Fisher Scientific, Cat. 11140-50), 0.1mM β-mercaptoethanol (Thermo Fisher Scientific, Cat. 21985-023) and Antibiotic-Antimycotic (Thermo Fisher Scientific, Cat. 15240-062). 2i inhibitors were also added into the medium at final concentrations of 1μM for PD0325901 (Millipore Sigma, Cat. PZ0162) and 3μM for CHIR99021 (STEMCELL Technologies, Cat. 72052), respectively.

### Generation of stable cell lines and genetic knock-in cells

We constructed plasmids PiggyBac-EF1-HaloTag-OCT4, PiggyBac-EF1-H2A.Z-HaloTag and PiggyBac-EF1-HaloTag-mTBP based on an available PiggyBac-EF1-HaloTag backbone vector. We used them together with the PiggyBac super-transposase to generate stable cell lines that express HaloTag-OCT4, H2A.Z-HaloTag and HaloTag-mTBP fusion proteins for single-particle tracking experiments. For H2B single-particle tracking experiment, we adopted a previously constructed PiggyBac-EF1-H2B-HaloTag (Li et al., 2016). Maps of plasmids used in this study are available upon request.

To generate the stable Halo-OCT4 and Halo-mTBP expressing mESCs, 8µg PiggyBac-EF1-HaloTag-OCT4 or PiggyBac-EF1-HaloTag-mTBP was nucleofected with 8µg PiggyBac super-transposase into ∼1×10^6^ mESCs. The derived mESCs were selected with 500µg/mL G418 (Thermo Fisher Scientific, Cat. 10131035) for ∼5 days, stained with JF549 dye and FACS-sorted for cell population with intermediately expressed genes.

We have applied previously described methods to generate sgRNA constructs and to construct Halo-MED1, MED6-Halo and Halo-BRD4 donor plasmids for generating knock-in cells by CRISPR/Cas9 genome editing technology (Hansen et al., 2017). The corresponding Emerald version of MED1/MED6 donor constructs were generated by replacing the HaloTag DNA coding sequence with Emerald coding sequence by Gibson assembly (NEB, Cat. E5520S). To generate Halo-MED1, MED6-Halo and Halo-BRD4 genetic knock-in cells, 1.0µg/µl SpCas9-sgRNA-PGK-Venus construct and 1µg/µl donor construct were nucleofected into ∼3×10^6^ Tir1 mESCs using the Amaxa™ 4D-Nucleofector and the P3 Primary Cell 4D-Nucleofector™ X Kit (Lonza, Cat. V4XP-3024) following the manufacture’s protocol. Three days following nucleofection, cells were then stained with 50nM Janelia Fluor^®^ 549 HaloTag ligand (JF549) for 30 mins, washed 3 times in 1×PBS and then in mESC medium for 15 mins, and subjected to FACS sorting. JF549-positive cells were plated sparsely in 60mm tissue culture plates and grown for another 5∼7 days. Single colonies were picked for genotyping by designing PCR primers outside of the homology arms. Biallelic knock-in mESC clones were verified by PCR genotyping, Sanger sequencing, and Western blot.

### Western Blots

mESCs were lysed in 1×SDS sampling buffer (200mM Tris HCl, pH=7, 10% glycerol, 2% SDS, 4% β-mercaptoethanol, 400mM DTT, 0.4% bromophenol blue). Lysates were sonicated and then incubated on ice for 30 mins, mixed with 2×loading buffer and denatured at 95°C for 5 mins. Lysed proteins were resolved by SDS-PAGE using Mini-PROTEAN^®^ TGX™ Precast Gels (Bio-Rad). Primary antibodies used include: anti-RAD21(D213) rabbit polyclonal antibody (Cell Signaling, Cat. 4321), anti-MED1(CRSP1/TRAP220) rabbit polyclonal antibody (Bethyl Laboratories Inc, Cat. A300-793A), anti-MED6 Rabbit polyclonal antibody (Abcam, Cat. ab220110), anti-BRD4 (E8V7I) Rabbit monoclonal antibody (Cell Signaling, Cat. 54615), anti-TBP Rabbit polyclonal antibody (Cell Signaling, Cat. 8515), anti-OCT4 rabbit polyclonal antibody (Abcam, Cat. ab19857), Anti-Histone H2A.Z antibody [EPR18090] (Abcam, Cat. ab188314) and anti-α-tubulin (11H10) rabbit monoclonal antibody (Cell Signaling, Cat. 2125). We used HRP-linked anti-rabbit IgG (Cell Signaling, Cat. 7074) secondary antibodies at a dilution of 1:1000. Western blots were exposed by using the 20×LumiGLO chemiluminescent detection system (Cell Signaling, Cat. 7003) and imaged by using a Bio-Rad ChemiDoc MP detection system.

### Cell cycle analysis

Cell cycle analysis was performed by propidium iodide (PI) staining following the protocols from the propidium iodide flow cytometry kit (Abcam, Cat. ab139418). Briefly, cells were trypsinized into single cell suspension and fixed with 67% ice-cold ethanol in PBS overnight at 4°C. The next day, the cells were rehydrated with PBS and then stained with propidium iodide (final concentration of 50μg/ml) and RNaseA (final 50μg/ml) for 30 mins at 37°C before flow cytometry analysis. All samples were acquired on a Beckman Coulter CytoFLEX S with 4 lasers (405nm, 488nm, 561nm, 638nm) (Beckman Coulter), and operated by CytExpert Software v2.3 (Beckman Coulter) using 488nm FSC-H (488-FSC-H) as the threshold parameter (threshold automatic setting). Detector’s gain for fluorescence and FSC/SSC detection were optimized by using control cells without treatment. The following gains were used for detectors with different spectral filters: 39, 15, 28, and 10 a.u. for 488-FSC, 488-SSC, 561-585/42 and 561-610/20 respectively. SSC-A versus FSC-W was used for initial gating of singlet cells followed by SSC-A versus FSC-A to further define cellular events. An event count versus 561-610/20 histogram plot was used to determine the percentage of cells in G1, S or G2/M phases of the cell cycle. All samples were acquired for 5 mins at a sampling rate of 30μl/min or up to 15,000 cells. Flowjo v.10.7.1 (Flowjo, LLC) was used for analysis of flow cytometry data.

### Smart-SCRB data acquisition

Mouse embryonic stem cells with engineered auxin-induced degron system treated with or without auxin were resuspended in culture medium without Phenol red, and FACS-sorted into 96-well PCR plates containing 3μl mild lysis buffer (nuclease-free water with 0.2% Triton + 0.1U/μl RNase inhibitor (Lucigen, Cat. 30281-2)). A total of ∼400 cells were sorted and collected for each sample. The PCR plates with collected cells were briefly centrifuged, immediately frozen and stored at -80 °C until cDNA synthesis.

1μl harsh lysis buffer (50mM Tris (pH=8.0), 5mM EDTA (pH=8.0), 10mM DTT, 1% Tween-20, 1% Triton X-100, 0.1g/l proteinase K, 2.5mM dNTPs, and ERCC Mix (10^7^-fold dilution) and 1μl 10mM barcoded RT primer was added to each well. Plates were incubated at 50°C for 5 mins to lyse cells and proteinase K was heat-inactivated by subsequently incubating at 80°C for 20 mins. To minimize contamination across wells, heavy-duty plate seals and qPCR compression pads (Thermo Fisher Scientific, Cat. 4312639) were used to seal the plates. The lysis reaction was mixed with 2μl reverse transcription master mix 5×buffer (Thermo Fisher Scientific, Cat. 11756500), 2μl 5M Betaine (Sigma-Aldrich, Cat. B0300-1VL), 0.2μl 50mM E5V6NEXT template switch oligo (Integrated DNA Technologies), 0.1μl 200U/ μl Maxima H-RT (Thermo Fisher Scientific, Cat. EP0751), 0.1μl 40U/μl NxGen RNase Inhibitor, and 0.6μl nuclease-free water (Thermo Fisher Scientific, Cat. AM9932). The reaction system was then incubated at 42°C for 1.5 hrs, followed by 10 mins at 75°C to inactivate reverse transcriptase. PCR was performed by adding 10μl 2×HiFi PCR mix (Kapa Biosystems, Cat. 7958927001) and 0.5μl 60mM SINGV6 primer and running the following program: 98°C for 3 mins, 20 cycles of 98°C for 20 s, 64°C for 15 s, 72°C for 4 mins, with a final extension step off 5 mins at 72°C.

Single-cell cDNA was pooled by plate to make library. 600 pg cDNA from each sample plate was used in a modified Nextera XT (Illumina, Cat. FC-131-1024) library preparation but using the P5NEXTPT5 primer and the tagmentation time of 5 mins. The resulting libraries were purified following the Nextera XT protocol (0.6× ratio) and quantified by qPCR using Kapa Library Quantification kit (Kapa Biosystems, Cat. KK4824). Four plates were pooled together on a NextSeq 550 high-output flow cell or NextSeq 2000 P2-100 flow cell with 26 bp in read 1 and 50 bp in read 2. PhiX control library (Illumina) was spiked in at a final concentration of 7.5% to improve color balance in Read 1. Read 1 contains the spacer, barcode, and UMI and read 2 represents a cDNA fragment from the 3’ end of the transcript. The entire experimental procedure was replicated two more times and ∼400 cells were analyzed for each condition (control vs. auxin-treated).

Alignment and count-based quantification of single-cell data was performed by removing adapters, tagging transcript reads to barcodes and UMIs, and aligning the resulting data to the mouse genome mm10. After quantification, 2,600 detected genes (gene_det > 2600) was set as a threshold to eliminate unreliable sequencing results and the data matrix (*M*_RNA_) for all remaining cells were used for analysis.

### Identification of ACDs

To identify ACDs across the genome, we referred to available bulk ATAC-seq data (Xie et al., 2022) and used 200kb genomic distance as the binning unit for analysis. The Matlab function findpeaks() was then called in “MinPeakProminence” mode and the value of “MinPeakProminence” was set to the average of overall ATAC signals. A total of 776 ACDs were identified by using these criteria.

### Gene-gene correlation analysis

Statistically significant gene correlations were calculated genome wide for all genes in which expression was detected (Figure S1D-E). UMI (Unique Molecular Identifier) data were used without normalization or log-transformation. Instead, corrected Pearson residuals were obtained for each gene and cell by subtracting from the observed UMI a gene- and cell-specific expectation value and dividing by the square root of that same value. Expectation values were determined by totaling the UMI for each gene, and, for each cell, dividing that total by the relative sequencing depth in that cell (total UMI by that cell as a fraction of total UMI in the entire dataset). Corrected Pearson residuals were further modified by dividing by the square root of 1+m/4, where m is the expectation value for that gene and cell. For any two genes, then, a modified, corrected Pearson correlation coefficient (mcPCC) was obtained by taking the dot product of vector of modified, corrected Pearson residuals, and dividing by the number of cells. This was then divided by the geometric mean of the average of the squared modified, corrected Pearson residuals for the two genes. By using modified, corrected Pearson residuals, this procedure corrects for unequal sequencing depth without adjusting the data matrix, and also corrects for overdispersion of highly expressed genes due to underlying stochasticity in their gene expression.

Because the statistical probability of observing any PCC varies markedly with the sparsity of the gene expression vectors used to calculate it, PCC magnitudes are not, themselves, a good measure of PCC significance. Therefore, significance was calculated directly, as follows: Using modified, corrected Pearson residuals, we construct the expected distribution of PCC values under the null hypothesis that each UMI is drawn from a Poisson Distribution whose mean is a random variate from a Lognormal distribution whose mean is the expectation value for that gene and cell, and whose coefficient of variation is ½. The first five moments of the distribution under the null hypothesis are then calculated exactly for each gene and cell, and from these the exact moments of the distributions of all 113,288,878 unique PCCs are determined by standard methods. Next, the Cornish Fisher approximation of the Edgeworth expansion1 is used to assign an approximate p-value to each observed mcPCC. To rigorously control for false discovery due to multiple hypothesis testing, we simulate the p-value distribution by constructing a matrix of simulated gene expression values for 5000 randomly selected genes, drawing for each cells a random variate from a Poisson Distribution whose mean is a random variate from a Lognormal distribution whose mean is the expectation value for that gene and cell, and whose coefficient of variation is ½. The identical procedure is applied to the simulated data set to produce a distribution of p-values. The observed and simulated p-value distributions are then compared to identify the largest p-value consistent with a false-discovery rate <0.05. Using this procedure, 841,635 significant correlations were identified, of which 46,916 involved pairs of genes on the same chromosome (19,384 positive and 27,532 negative).

The following procedure was used to determine whether intra-ACD correlations were more frequent than other intra-chromosomal correlations. First, we divided the number of correlations among genes in the same ACD by the number of expressed genes in that ACD to produce an observed correlation density. As the mean number of expressed genes per ACD was 5 ± 0.25, we then calculated the correlation density within sliding windows of 6 contiguous genes across each chromosome (use of alternate window sizes between 3 and 50 produced similar results). The results were plotted as a histogram displaying the frequency with which different correlation densities were observed, with ACD and genome wide, for positive and negative correlations separately.

### Gene co-expression analysis

To evaluate Spearman’s rank-order correlation among gene pairs, we filtered out low-expressed genes from *M*_RNA_ with a threshold of 1 after averaging the RNA-seq counts over the single cells analyzed. For each pair of expressed genes, the Spearman correlation coefficient *S*_x,y_ (considering tied ranks) was calculated by

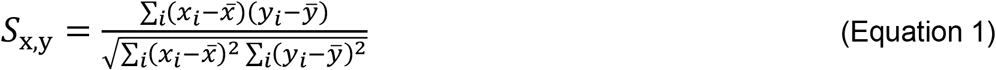

*x*_*i*_ and *y*_*i*_ are the sequencing counts for gene *x* and *y* in *i*^th^ individual cell, respectively. For each ACD pair – ACD1 and ACD2, the Spearman correlation matrix *S* was determined by calculating the Spearman correlation coefficient for each gene pair from ACD1 and ACD2, and the average Spearman correlation coefficient 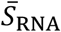 was calculated by taking the average value across the whole matrix *S* for control and Cohesin-depleted conditions.

The differential Spearman correlation matrix *ΔS*_RNA_ was calculated by

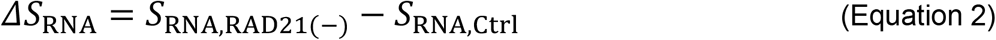

where *S*_RNA,RAD21(-)_ and *S*_RNA,Ctrl_ are Spearman correlation matrices for Cohesin-depletion and control conditions, respectively. The workflow for the above analyses is illustrated in Figure S1F.

The gene co-expression coefficient Δ*C*_*R*NA_(*i*, *j*)between two ACDs within the same chromatin was estimated by the following equation

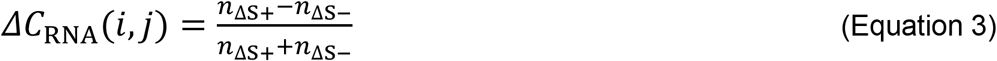

where *n*_ΔS+_ and *n*_ΔS−_ represent the number of positive and negative elements within the differential Spearman correlation matrix *ΔS*_RNA_. The derived differential gene co-expression matrix *ΔC*_RNA_ was plotted as a heatmap. The workflow for evaluating *ΔC*_RNA_ is illustrated in Figure S1G.

### Cell cycle phase classification based on Smart-SCRB data

To dissect the effect of cell cycle stages on gene co-expression analysis, we adopted a computational method described by Scialdone et al. for classifying cells into cell cycle phases based on Smart-SCRB data (Scialdone et al., 2015). Using a reference dataset, the difference in expression between each pair of genes was computed. Pairs with significant changes across cell cycle phases are selected as markers for classification and applied to a test dataset. Cells can then be classified into the appropriate phase, based on whether the observed change for each marker pair is consistent with one phase or another. This approach is implemented in the *cyclone()* function from the “*scran*” package, which contains pre-trained set of marker pairs (mouse_cycle_markers.rds) for mouse cells.

### Single-cell ATAC-seq data acquisition

Mouse embryonic stem cells treated with or without auxin were washed and resuspended in 1xPBS with 0.04% BSA. Nuclei isolation for single-cell ATAC (scATAC) sequencing from cell suspensions was carried out according to the manufacturer’s demonstrated protocol (Demonstrated Protocol, CG000169, Rev E,10xGenomics). Nuclei were counted using a Luna-II automated cell counter (Logos Biosystems). Approximately 15,000 nuclei per sample were loaded and subjected to the Chromium NextGem scATAC-seq v2 assay (10X Genomics). The resulting libraries were sequenced on a NextSeq 2000 (Illumina) with 50bp read 1, 8bp i7 index read, 16bp i5 index read, and 50bp read 2.

FASTQ files of raw data were processed by Cell Ranger ATAC (10xGenomics, v2.1.0) analysis pipeline. Reads were filtered and aligned to mouse genome mm10 (10xGenomics, refdata-cellranger-arc-mm10-2020-A-2.0.0) using *cellranger-atac count()* function with default parameters. Then the barcoded and aligned fragment files were loaded by ArchR (version 1.0.1). Low quality cells with minimum TSS enrichment score less than 4 and minimum fragment number less than 1000 were filtered out. Doublets were inferred by *addDoubletScores()* function and removed using *filterDoublets()* function with default parameters.

### Cross-ACD chromatin co-accessibility analysis

To evaluate the chromatin co-accessibility per ACD pair, we binned ATAC counts within each ACD to derive an ACD (*M*_ATAC_) for each experimental condition. After adding ACDs regions by *addPeakSet()* function, counts for each ACD per cell were aggregated together by *addPeakMatrix()* with a very high ceiling value (1000). The ACD counts matrix (*M*_ATAC_) for each condition was normalized to the average value 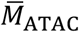 of the matrix to derive a normalized ACD counts matrix (*N*_ATAC_). To mitigate the influence from poorly sequenced cells (have 0 count for many ACDs), each column *j* of *N*_ATAC_ with the mean value 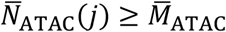 was selected and integrated to form a new matrix *N*_ATAC-Filtered_ for Spearman’s rank-order correlation evaluation.

For each ACD pair, the Spearman correlation coefficient for chromatin co-accessibility was calculated by using Equation 1. The Spearman correlation matrix *S*_ATAC_ was determined by calculating the Spearman correlation coefficient for each ACD pair across the genome. The differential Spearman correlation matrix *ΔS*_ATAC_ was calculated by

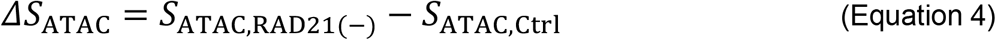

where *S*_ATAC,RAD21(-)_ and *S*_ATAC,Ctrl_ are Spearman correlation matrices for Cohesin-depletion and control conditions, respectively. *S*_ATAC,RAD21(-)_, *S*_ATAC,Ctrl_ and *ΔS*_ATAC_ were plotted as heatmaps. The workflow for evaluating Spearman correlation coefficient for chromatin co-accessibility is illustrated in Figure S2A.

### Single-molecule RNA-FISH

Single-molecule (sm) RNA-FISH probe blends were designed through Stellaris RNA FISH probe designer (Biosearch Technologies) and include 40∼48 serial probes which target only the intron regions of selected genes. The probe pairs designed for gene pairs selected from neighbor TADs were labelled by Quasar 570 and Quasar 670 respectively for two-color experiments. The commercially synthesized oligonucleotide probe blends were dissolved in 400μl of TE buffer (10mM Tris-HCl, 1mM EDTA, pH=8.0) to create a probe stock of 12.5μM.

smRNA-FISH experiments were performed according to the Stellaris^®^ RNA FISH protocol for adherent cells provided by Biosearch Technologies. Specifically, cells were grown on 18mm round #1 cover glass (Warner Instruments, Cat. CS-18R) in a 12-well cell culture plate coated with Human recombinant laminin 511 (BioLamina, Cat. LN511-0202) and fixed in 1xPBS with 3.7% formaldehyde (Millipore Sigma, Cat. F8775-25ML) for 10 mins and then permeabilized in ice-cold 70% (vol./vol.) ethanol for 2 hrs. The coverslips with cells were immersed in 100µl hybridization buffer (90µl Stellaris RNA-FISH Hybridization Buffer (Biosearch Technologies, Cat. SMF-HB1-10) and 10µl deionized formamide (Millipore Sigma, Cat. S4117)) at a final probe concentration of 125nM and placed into a humidified chamber. The assembled humidified chamber was incubated overnight at 37°C, before the sample coverslips were washed, co-stained with 5ng/ml DAPI (Sigma-Aldrich, Cat. D8417) and mounted onto slides for future imaging analysis.

### Chr 2 Intron-FISH probe design, synthesis and amplification

For 208 active genes across mouse chromosome 2, we selected non-overlapping 35-nt probes with several constraints including max TM 100°C, min TM 74°C, <u>secondary structure</u> temp 76°C, cross hyb temp 72°C, 30%–90% <u>GC content</u>, no more than 6 contiguous identical <u>nucleotides</u> and at least 2-nt spaces between adjacent probes. Primary probes were screened for potential non-specific binding with Bowtie2 (--very-sensitive-local) against mm10 genome sequences. Probes with more than one binding sites were filtered out. 30 qualified probes per gene closest to the transcription start site were selected. 3 nt spacer (random sequence) were extended at the 5′ and 3’ end of the 35-nt probe sets. 2 readout sequences (15nt) separated by a 2nt spacer (random sequence) were added at both 5’ and 3’ of the probe respectively for the potential of performing sequential FISH experiments. Then, universal primer sequences were attached at 5′ and 3′ ends. The 5’ primer contains a T7 promoter. The total length of each probe is 147nt. The <u>oligonucleotide probe</u>pool (6180 probes) was purchased from Twist Bioscience.

For probe amplification, limited PCR cycles were used to amplify the designated probe sequences from the oligo complex pool with kapa HiFi HotStart Polymerase (Roche, Cat. KK2502). Then, the amplified PCR products were purified using the Zymo dna clean and concentrator kit (Zymo Research, Cat. D4014) according to the manufacturer’s instructions. The PCR products were used as the template for in *vitro* transcription (NEB, Cat. E2040S) followed by reverse transcription (Thermo Fisher Scientific, Cat. EP7051) with the forward primer. After reverse transcription, the probes were subjected to uracil-specific excision reagent (USER) enzyme (NEB, Cat. N5505S) treatment for ∼24 h at 37 °C. Then, probes were alkaline hydrolysed with 1M NaOH at 65 °C for 15 mins to degrade the RNA templates, followed by 1M acetic acid neutralization. Next, to clean up the probes, we used ssDNA/RNA Clean & Concentrator (Zymo Research, Cat. D7011) before hybridization.

### Chr 2 Intron-FISH

The Chr2 Intron-FISH was performed following a revised seq-FISH protocol as described by Shah et al. (Shah et al., 2018). Mouse embryonic stem cells were plated on Human recombinant laminin 511-coated coverslips (Electron Microscopy Sciences, Cat. 72196-25). Cells were then fixed using 4% formaldehyde (Thermo Fisher Scientific, Cat. 28908) in 1×PBS diluted in molecular biology grade water (Corning, Cat. 46-000-CM) for 15 min at 20°C, washed with 1×PBS for a few times, and incubated in 70% ethanol for about 3 hours at room temperature.

The coverslips were then washed twice with 2×SSC. For primary probe hybridization, samples were incubated with primary Chr 2 intron probes for 30 hrs at 37°C in 50% Hybridization Buffer (2×SSC, 50% Formamide (v/v) (Thermo Fisher Scientific, Cat. AM9344), 10%(w/v) Dextran Sulfate (Sigma-Aldrich, Cat. D8906) in molecular biology grade water), and then washed in 55% Wash Buffer (2× SSC, 55% Formamide (v/v), 0.1% Triton X-100 (Sigma-Aldrich, Cat. 93443)) for 30 mins at room temperature, followed by 2×SSC wash. The Alexa 647-coupled Imager probe (Integrated DNA Technologies) for the first round of hybridization were incubated for 20 mins at 50 nM each at room temperature in 10% EC buffer (10% ethylene carbonate (Sigma-Aldrich, Cat. E26258), 2×SSC, 0.1g/ml Dextran Sulfate and 0.02U/ml SUPERase·In™ RNase Inhibitor (Thermo Fisher Scientific, Cat. AM2694)) and washed for 5 mins at room temperature in 10% Wash Buffer (2×SSC, 10% formamide (v/v), 0.1% Triton X-100) followed by 1 min wash in 2×SSC.

Samples were then co-stained with 5ng/ml DAPI for 15 mins and imaged in an anti-bleaching buffer (50mM Tris-HCl, pH=8.0, 300mM NaCl, 2×SSC, 3mM Trolox (Sigma-Aldrich, Cat. 238813), 0.8% D-glucose (Sigma-Aldrich, Cat. G7528), 100-fold diluted Catalase (Sigma-Aldrich, Cat. C3155), 0.5mg/ml Glucose Oxidase (Sigma-Aldrich, Cat. G2133) and 0.02U/ml SUPERase·In™ RNase Inhibitor).

### Airyscan imaging and image analysis

Fluorescence images obtained from smRNA-FISH or Chr 2 Intron-FISH were acquired on the ZEISS LSM880 Inverted Confocal microscope attached with an Airyscan 32 GaAsP (gallium arsenide phosphide)-PMT area detector. Before imaging, the beam position was calibrated to center on the 32-detector array. Images were taken under the Airyscan super-resolution mode by a Plan Apochromat 63×/NA1.40 oil objective in a lens immersion medium with a refractive index of 1.515. The Airyscan super-resolution technology used a very small pinhole (0.2AU) at each of its 32 detector elements to increase SNR ∼4-8 fold and enables ∼1.7-fold improvement of resolution upon linear deconvolution in both lateral (x, y) and axial (z) directions. The DAPI, mEmerald, Quasar 570 (and JF549) and Quasar 670 signals were illuminated/detected at excitation/emission wavelength of 405nm/460nm, 488nm/510nm, 561nm/594nm and 633nm/654nm, respectively. Z-stacks were acquired with a step of 300nm. After image acquisition, Airyscan image was processed and reconstructed using the provided algorithm from ZEISS LSM880 platform.

3D Airyscan image stacks were processed by using Imaris 7.2.3. As an initial step, we manually inspected the images and removed the low-quality images based on the following criteria: (1) non-specific signal outside of the DAPI-stained nuclei, (2) cropped signal at the edge of the images, (3) very faint signal.

To characterize and visualize MED6 and MED1 protein hubs, we employed the Surfaces object in Imaris and ran the following algorithms: (1) apply “Background Subtraction (Local Contrast)” mode and set “Diameter of Largest Sphere” as 800 nm; (2) uniformly use threshold value 30 for MED6 and 10 for MED1 to segment protein hubs; (3) set the “Minimal Number of Voxels” as 20 for filtering out noise signals. To characterize and visualize smRNA-FISH puncta, we employed the Surfaces object from Imaris and and ran the following algorithms: (1) apply “Background Subtraction (Local Contrast)” mode and set “Diameter of Largest Sphere” as 2000 nm; (2) uniformly use threshold value 35 for Quasar 570 and 70 for Quasar 670 to illustrate smRNA-FISH puncta; (3) set the “Minimal Number of Voxels” as 50 for filtering out noise signals. To characterize and the colocalization signal between Mediator hubs and smRNA-FISH puncta, we employed the “Colocalization” function in Imaris with a common threshold value of 100 to identify the overlapping volumes and reconstruct the 3D *iso*-surfaces.

To measure the 3D distance between puncta of different smRNA-FISH signals, we localized the voxels corresponding to the local maximum of identified RNA FISH signal using the Imaris “Spots” function module and calculated the Euclidean distance by using Measurement Points object with the voxel size (43.6nm × 43.6nm × 300nm). The potential drifts among different imaging channels were estimated by using 100nm multi-spectral beads under the same acquisition settings and was considered into the calculation of Euclidean distance between smRNA-FISH foci. The gene-pair smRNA-FISH data was screened for a number (*N*) of cells. The number of bursting (*N*_b_) events for each gene was counted when a visible smRNA-FISH puncta was identified. The number of co-bursting (*N*_co_) events for each gene pair was counted when a visible and approximated (Euclidean distance < 2µm) pair of smRNA-FISH puncta from two channels were identified. The bursting fraction (*F*) of each gene was computed by

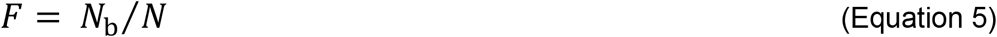

The co-bursting fraction (*F*_co_) of gene pair (A and B) was computed by

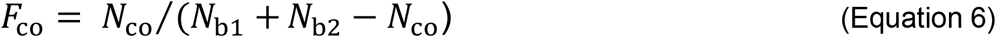

*N*_b1_ and *N*_b2_ are the numbers of bursting events for A and B, respectively.

### Gene co-expression simulation

The Gillespie algorithm (Gillespie, 1977) allows the modelling of exact dynamics described by the master equation in single cells. We applied this algorithm to simulate the mRNA and protein expression dynamics for Gene A and B. A stoichiometry matrix (number of reactions × number of reactants) was constructed with parameter values describing the basic reaction steps – transcription, translation, mRNA degradation and protein degradation (Figure S4A). To simulate the expression dynamics of A and B without co-regulation, transcription of A and that of B were formulated as independent reaction steps; In contrast and to simulate the expression dynamics of A and B with co-regulation, a co-regulation step for both A and B was introduced into the stoichiometry matrix. Simulations were performed with parameter values set within the range of physiological level. For each condition, 5,000 trajectories (corresponding to 5,000 cells) were simulated and the results at the end point was analyzed and plotted. Sensitivity analysis on Spearman’s correlation coefficient with or without co-bursting was done by perturbing each of the parameter values within the range of 0.1∼10 fold.

For the numerical simulation of co-expression involving multiple gene pairs, 10,000 cells were randomly assigned to co-express each pair based on the co-expression frequency. Cells that carry at least one co-activated gene pair were counted and used for the fraction calculation.

### 3D ATAC-PALM, Mediator-HaloTag PALM and image analysis

We prepared the reagents for 3D ATAC-PALM experiments as described previously (Xie et al., 2020). One day before experiment, cells were plated on 5mm coverslips (Warner Instruments, 64-0700) at around 70-80% confluency with proper coating. Cells were fixed with 3.7% formaldehyde (Millipore Sigma, Cat. F8775-25ML) for 10 mins at room temperature. After fixation, cells were washed three times with 1×PBS for 5 mins and then permeabilized with ATAC lysis buffer (10mM Tris-Cl, pH=7.4, 10mM NaCl, 3mM MgCl_2_, 0.1% Igepal CA-630) for 10 mins at room temperature. After permeabilization, the sample coverslips were washed twice in 1×PBS and the transposase mixture solution (1× Tagmentation buffer-10mM Tris-HCl, pH=7.6, 5mM MgCl_2_, 10% dimethylformamide, 100 nM Tn5-PA-JF_549_) was then added to the sample. The coverslips were placed in a humidity chamber and incubated for 30 mins at 37 °C. After the transposase reaction, the coverslips were washed three times in 1×PBS containing 0.01% SDS and 50mM EDTA for 15 mins at 55°C before mounted onto the lattice light-sheet microscope (LLSM) sample stage for imaging.

The 3D ATAC-PALM data were acquired by the lattice light-sheet microscopy at room temperature (Chen et al., 2014a). The light sheet was generated from the interference of highly parallel beams in a square lattice and dithered to create a uniform excitation sheet. The inner and outer numerical apertures of the excitation sheet were set to be 0.44 and 0.55, respectively. A Variable-Flow Peristaltic Pump (Fisher Scientific, Cat. 13-876-1) was used to connect a two-liter reservoir with the imaging chamber with 1×PBS circulating through at a constant flow rate. Labelled cells seeded on 5mm coverslips were placed into the imaging chamber and each imaging volume took 100∼200 image frames, depending on the depth of the field of view. Specifically, spontaneously activated PA-JF_549_ dye were initially pushed into the fluorescent dark state through repeated photo-bleaching by scanning the whole imaging volume with a 2W 560nm (or 640nm) laser (MPB Communications Inc., Canada). Then, the samples were imaged by iteratively photo-activating each plane with very low-intensity 405nm light (<0.05mW power at the rear aperture of the excitation objective and 6W/cm^2^ power at the sample) for 8 ms and by alternatively exciting each plane with a 2W 560nm laser and a 2W 640nm laser at its full power (26mW power at the rear aperture of the excitation objective and 3466W/cm^2^ power at the sample) for 20 ms exposure time. The specimen was illuminated when laser light went through a custom 0.65 NA excitation objective (Special Optics, Wharton, NJ) and the fluorescence generated within the specimen was collected by a detection objective (CFI Apo LWD 25×W, 1.1 NA, Nikon), filtered through a 440/521/607/700nm BrightLine quad-band bandpass filter (Semrock) and N-BK7 Mounted Plano-Convex Round cylindrical lens (f=1000mm, Ø 1”, Thorlabs, Cat. LJ1516RM), and eventually recorded by an ORCA-Flash 4.0 sCMOS camera (Hamamatsu, Cat. C13440-20CU). The cells were imaged under sample scanning mode and the dithered light sheet at 500 nm step size, thereby capturing a volume of ∼25µm × 51µm × (27∼54) µm, considering 32.8° angle between the excitation direction and the stage moving plane.

To precisely analyze the 3D ATAC-PALM data, we embedded nano-gold fiducials within the coverslips for drift correction as previously described (Xie et al., 2020). ATAC-PALM images were taken to construct a 3D volume when the sample was moving along the “s” axis. Individual volumes per acquisition were automatically stored as Tif stacks, which were then analyzed by in-house developed scripts in MATLAB. The cylindrical lens introduced astigmatism in the detection path and recorded each isolated single molecule with its ellipticity, thereby encoding the 3D position of each molecule relative to the microscope focal plane. All processing was performed by converting all dimensions to units of x-y pixels, which were 100nm×100nm after transformation due to the magnification of the detection objective and tube lens. We estimated the localization precision by calculating the standard deviation of all the localizations coordinates (x, y and z) after the nano-gold fiducial correction. The localization precision is 26±3nm and 53±5 nm for xy and z, respectively.

### 3D pair cross-correlation function

3D pair cross-correlation function *c*_0_(*r*) between localizations of molecule A and those of molecule B can be formulated as

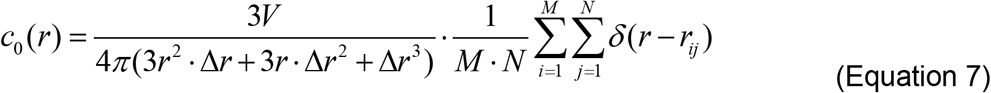

*M* is the total number of localizations for molecule A and *N* is the total number of localizations for molecule B. Δ*r* = 50nm is the binning width used in the analysis. The Dirac Delta function is defined by

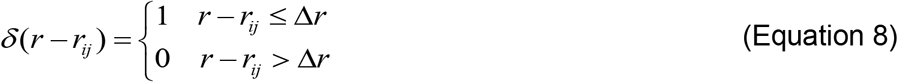

Where *r*_*ij*_ represents the pair-wise Euclidean distance between localization point *i* and *j*. The normalized 3D pair auto-correlation function *C*(*r*) was calculated by:

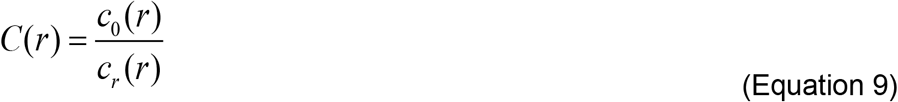

*c*_*r*_(*r*) refers to the pair cross-correlation function calculated from uniform distributions with the same localization density in the same volume as real data used.

### Single-molecule imaging

Single-molecule imaging experiments were carried out as previously described (Chen et al., 2014b; Lerner et al., 2020) on a Nikon Eclipse TiE Motorized Inverted microscope equipped with a 100× Oil-immersion Objective lens (Nikon, N.A.=1.49), four laser lines (405/488/561/642nm), an automatic TIRF illuminator, a perfect focusing system, a tri-cam splitter, three EMCCDs (iXon Ultra 897, Andor) and Tokai Hit environmental control (humidity, 37°C, 5% CO_2_). Proper emission filters (Semrock) were switched in front of the cameras for GFP and JF549 emission and a band mirror (405/488/561/633nm BrightLine quad-band bandpass filter, Semrock) was used to reflect the laser into the objective.

To perform single-molecule imaging of TFs and cofactors, cells were seeded on 25mm #1.5 coverglass pre-cleaned with potassium hydroxide (KOH) and ethanol and coated with Human recombinant laminin 511 according to manufacturer’s instruction. Live cell imaging experiments were conducted by culturing mESCs in imaging medium composed of FluoroBrite DMEM (Thermo Fisher Scientific, Cat. A1896701), 10% FBS, 1×GlutaMax, 1×NEAA, 0.1mM β-mercaptoethanol and 1000U/ml LIF. The TIRF illuminator was adjusted to deliver a highly inclined laser beam to the cover glass with the incident angle smaller than the critical angle. The oblique illumination (HILO) has much less out-of-focus excitation, compared with the regular Epi-illumination. TFs and cofactors linked to HaloTag were labeled with 5nM Halo ligand-JF549 for 15 mins and imaged using a 561 nm laser with the excitation intensity of ∼50 W/cm^2^. To minimize drift, the imaging experiments were performed in an ultra-clear room with the precise temperature control system. The environment control chamber for cell culturing was thermo-equilibrated. The imaging system was calibrated with beads to confirm a minimal drift during imaging (x-y drift < 100nm per hour).

For fast-molecule tracking and throughout the experiments, we used 10 ms imaging acquisition time and took 5,000 continuous frames per imaging view after photo-bleaching of saturatedly labelled single molecules at the beginning. About 10∼15 views were imaged for each labelled transcription factor/cofactor under normal or Cohesin-depleted conditions. In contrast for evaluating the radius of confinement which captures the motility of stably bound molecules, we used a longer acquisition time of 50 ms and took 5,000 continuous frames per imaging view.

### Single particle tracking analysis

Each imaging view was recorded as a TIF stack and single molecules were tracked using SLIMfast, a custom-written MATLAB implementation of the MTT algorithm (Serge et al., 2008). Frame-to-frame motions are defined by the distance between consecutive positions of the particle and can be potentially related to (a) Brownian (random walk) or confined motions of the molecule (b) potential artifactual effects such as imperceptible movements of the nucleus. To filter out the effects resulting from (b), it is thus necessary to define a Maximal Expected Diffusion Coefficient (*D*_Max_), that defines the maximal distance (*d*_m_) between two consecutive frames for a particle to be considered as the same object. As in the previous publication (Chen et al., 2014b), a cutoff was set to 3*d*_m_ to ensure a 99% confidence level and *D*_Max_ was set as 1μm^2^/s^-1^.

For each imaging view, SLIMfast generated an .txt output file consisting of a series of successive x/y coordinates and times of detection, corresponding to the displacement of each individual molecule. The output SLIMfast .txt files include the following information: x/y coordinates (2D coordinates of the molecule in μm), trajectory index (ID number of the trajectory) and frame number (the index of the frame on which each single molecule was detected). These track files were used as the inputs to perform two-state (for histone subunits) or three-state (for transcriptional regulators) kinetic fitting by using Spot-On (Hansen et al., 2018) software in order to compute the biophysical parameter values of single particles.

The radius of confinement (*RC*) represents the circle best encompassing the motion track, rather than encompassing it strictly. Thus, the measurement of the radius of confinement is largely independent of the track duration. Tracks with length<5 were discarded in the preprocessing step. To quantify *RC*, the mean square displacement (MSD) curves of each track were fitted using nonlinear least-squares approach in Matlab with a circle confined diffusion model (Wieser and Schutz, 2008) as illustrated in the following equation:

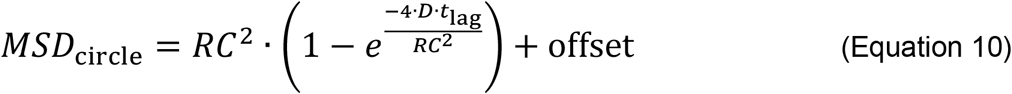

The fitting provided values for *RC*, the diffusion coefficient at short timescales *D*, and a constant offset value due to the localization precision limit which is inherent to all the localization-based microscopy methods. To discard fitting errors related to artifacts such as erroneously connected jumps, we have discarded the trajectories with squared norm of the residual higher than 10^−5^, and *RC* higher than 500nm.

### Statistical Analysis

Unless specified, data are presented as Mean ± Standard Deviation (SD) with statistical significance (* p < 0.05, ** p < 0.01, *** p <0.001). We normally applied Student’s *t*-test and non-parametric Wilcoxon test as specified in the figure legends.

## Acknowledgments

We thank the Tjian-Darzacq lab members for helpful discussions and constructive suggestions. We also thank Kathy Schaefer from Cell and Molecular Biology Shared Resources in Janelia Research Campus (JRC) for assistance in cell sorting and CytoFLEX flow cytometry experiments, Michael DeSantis and Damien Alcor from Imaging Core facility at Janelia for microscopy training, and Melanie Radcliff for administrative support.

## Funding

P.D., L.X., L.W., A.L.L. and Z.J.L. are funded by the Howard Hughes Medical Institute (HHMI). L.X. also receives support from the Janelia Visitor Program. S.Z. and H.Y.C. are funded by HHMI and NIH RM1-HG007735 and R35-CA209919. A.D.L was supported by NIH grants U54-CA217378 and R01-DE019638.

## Author contributions

P.D. and Z.J.L. conceived and designed the study. P.D. constructed reagents/cell lines, performed imaging (3D ATAC-PALM, Airyscan and single molecule tracking), biochemical experiments and data analysis. P.D. and S.Z. performed Smart-SCRB and scATAC-seq analysis. L.X. helped with cell line construction. L.W., and A.L.L. performed Smart-SCRB and scATAC-seq experiments of samples provided by P.D.

A.D.L. conducted the gene-gene correlation analysis on Smart-SCRB data. P.D. and Z.J.L. wrote the manuscript with inputs from other authors. Z.J.L. and H.Y.C. supervised the study.

## Declaration of interests

Authors declare no competing interests.

## Disclosure

H.Y.C. is a co-founder of Accent Therapeutics, Boundless Bio, Cartography Biosciences, Orbital Therapeutics, and is an advisor of 10xGenomics, Arsenal Biosciences, and Spring Discovery.

