## Supplementary Information for "Spatial organization of the 3D genome encodes gene co-expression programs in single cells"

---

Supplementary Figures ........................................ Page 2-16

Supplementary Movie Legends ........................................ Page 17

Supplementary Table Legends ........................................ Page 17

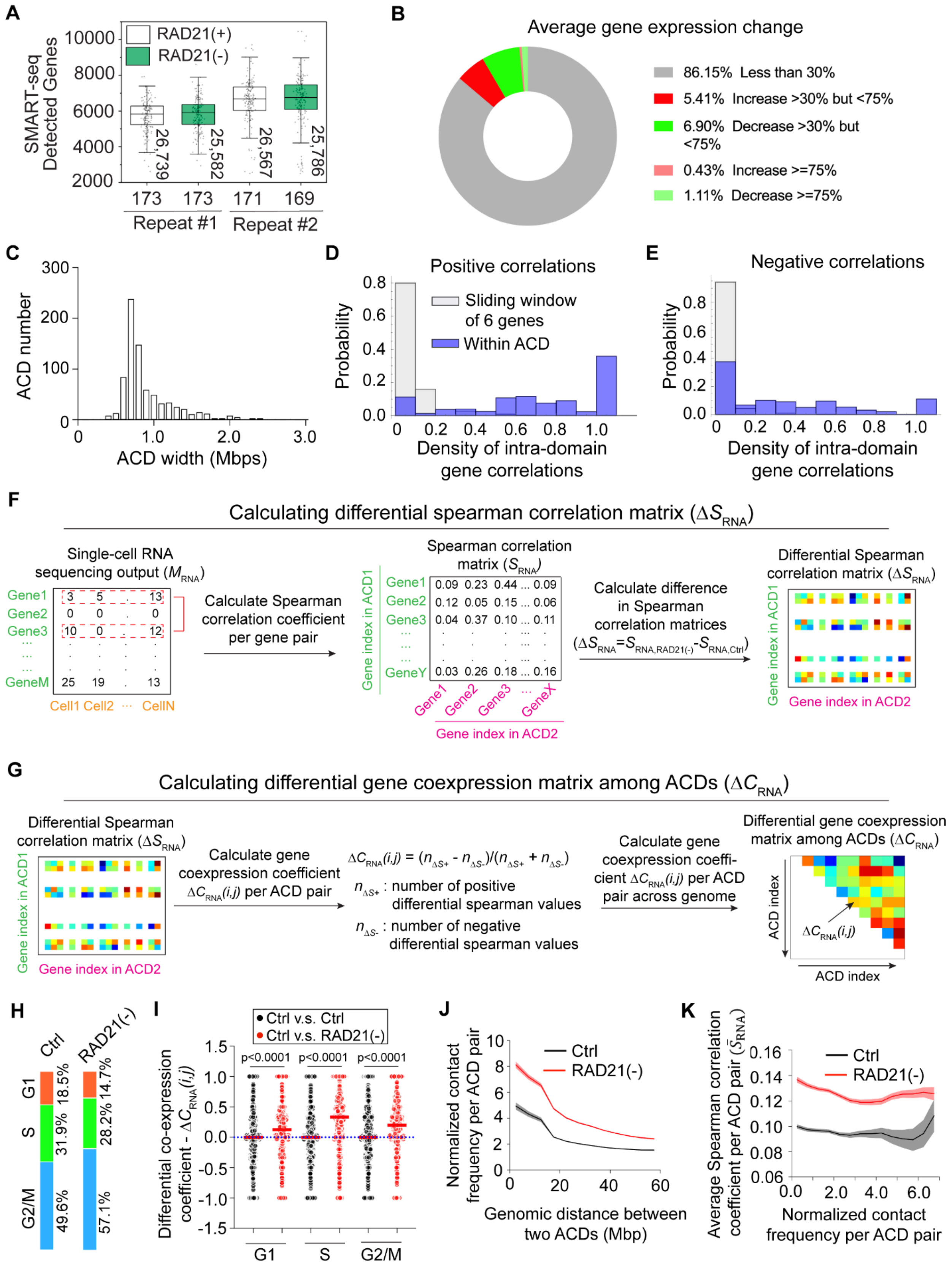

A
SMART-seq Detected Genes
10000
8000
6000
4000
2000
RAD21(+)
RAD21(-)
26,739
25,582
26,567
25,786
173
173
171
169
Repeat #1
Repeat #2
B
Average gene expression change
86.15% Less than 30%
5.41% Increase >30% but <75%
6.90% Decrease >30% but <75%
0.43% Increase >=75%
1.11% Decrease >=75%
C
ACD number
ACD width (Mbps)
D
Positive correlations
Sliding window of 6 genes
Within ACD
Probability
Density of intra-domain gene correlations
E
Negative correlations
Probability
Density of intra-domain gene correlations
F
Calculating differential spearman correlation matrix (ΔS_RNA)
Single-cell RNA sequencing output (M_RNA)
Gene1
Gene2
Gene3
GeneM
Cell1 Cell2 ··· CellN
Calculate Spearman correlation coefficient per gene pair
Spearman correlation matrix (S_RNA)
Gene index in ACD1
Gene1
Gene2
Gene3
GeneY
0.09 0.23 0.44 ... 0.09
0.12 0.05 0.15 ... 0.06
0.04 0.37 0.10 ... 0.11
0.03 0.26 0.18 ... 0.16
Gene1 Gene2 Gene3 ... GeneX
Gene index in ACD2
Calculate difference in Spearman correlation matrices
(ΔS_RNA=S_RNA,RAD21(-)-S_RNA,Ctrl)
Differential Spearman correlation matrix (ΔS_RNA)
Gene index in ACD1
Gene index in ACD2
G
Calculating differential gene coexpression matrix among ACDs (ΔC_RNA)
Differential Spearman correlation matrix (ΔS_RNA)
Gene index in ACD1
Gene index in ACD2
Calculate gene coexpression coefficient ΔC_RNA(i,j) per ACD pair
ΔC_RNA(i,j) = (n_ΔS+ - n_ΔS-)/(n_ΔS+ + n_ΔS-)
n_ΔS+ : number of positive differential spearman values
n_ΔS- : number of negative differential spearman values
Calculate gene coexpression coefficient ΔC_RNA(i,j) per ACD pair across genome
Differential gene coexpression matrix among ACDs (ΔC_RNA)
ACD index
ΔC_RNA(i,j)
ACD index
H
Ctrl
RAD21(-)
G1
S
G2/M
18.5%
31.9%
49.6%
14.7%
28.2%
57.1%
I
Ctrl v.s. Ctrl
Ctrl v.s. RAD21(-)
p<0.0001
p<0.0001
p<0.0001
Differential co-expression coefficient - ΔC_RNA(i,j)
G1
S
G2/M
J
Ctrl
RAD21(-)
Normalized contact frequency per ACD pair
Genomic distance between two ACDs (Mbp)
K
Ctrl
RAD21(-)
Average Spearman correlation coefficient per ACD pair (S̄_RNA)
Normalized contact frequency per ACD pair

**Figure S1. Quantification of cross-domain gene co-expression.**

**(A)** Box and dot plots of the number of detected genes per cell under control and Cohesin-depletion conditions for two biological replicates. The total number of genes detected across the population is annotated vertically. For all box charts, upper and lower whiskers represent outlier cut-offs based on the 1.5 interquartile range rule; the box represents the range from 25% to 75% percentile; the center line represents the median.

**(B)** The pie plot shows the degrees of global gene expression changes from pooled Smart-SCRB data.

**(C)** The size distribution of 776 ACDs across the mouse genome.

**(D-E)** Histograms show that both positive (**D**) and negative (**E**) correlations for gene co-expression are enriched within ACDs. X axis is the number of intra-ACD co-expression gene pairs divided by the number of detected genes within that domain. ACDs with no detected genes were excluded. As the control (grey bars), all contiguous detected genes were scanned for correlations in a sliding window of groups of 6 genes (about the median number for detected genes per ACD).

**(F)** The workflow for calculating differential Spearman correlation matrix ($\Delta S_{\mathrm{RNA}}$) per gene pair (from two ACDs) before and after Cohesin depletion from single-cell RNA-seq count matrix ($M_{\mathrm{RNA}}$).

(**G**) The workflow for calculating differential gene co-expression matrix ($\Delta C_{\mathrm{RNA}}$) per ACD pair before and after Cohesin depletion from $\Delta S$ in (**F**).

**(H)** Computational assignment of single cells into their cell cycle phases by analyzing Smart-SCRB data with *cyclone()* (R-programmed cell-cycle phase classifier.

**(I)** Dot plots show the distributions of differential gene co-expression coefficients before and after Cohesin depletion (red dots) for cells in G1, S and G2M cell cycle phases, respectively. Differential gene co-expression coefficients between two independent control groups (black dots) were used as the control. Red lines indicate the median values, and the blue dotted line indicates the zero-change line. Non-parametric Wilcoxon test was used for statistical testing.

**(J)** Normalized contact frequency per ACD pair (from Hi-C data) as a function of genomic distance between that ACD pair before and after Cohesin depletion. The normalized Hi-C contact frequency per ACD pair was calculated by averaging the frequencies of contacted regions within these two ACDs and plotted as a function of the genomic distance between that ACD pair after a five-point smoothing. Only the data from ACD pairs within the same chromosome were used to generate the plot. Shadow regions indicate standard error (S.E.) of the curves.

**(K)** Average Spearman correlation coefficient ($\bar{S}_{\mathrm{RNA}}$) per ACD pair as a function of normalized Hi-C contact frequency per ACD pair before and after Cohesin depletion. $\bar{S}_{\mathrm{RNA}}$ was plotted as a function of normalized contact frequency per ACD pair after a five-point smoothing. Shadow regions indicate standard error (S.E.) of the curves.

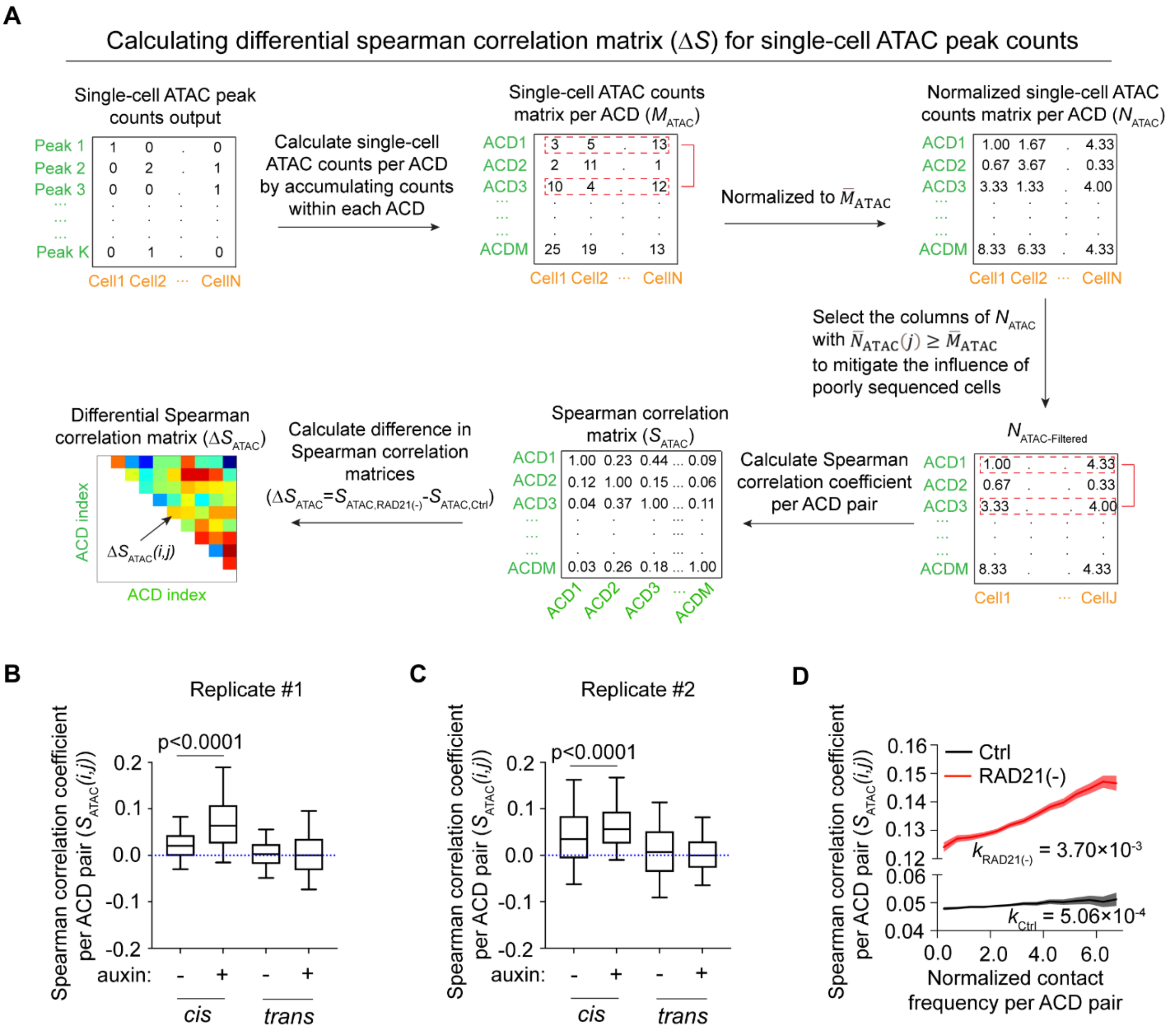
A
Calculating differential spearman correlation matrix (ΔS) for single-cell ATAC peak counts
Single-cell ATAC peak counts output
Peak 1
Peak 2
Peak 3
Peak K
Cell1 Cell2 ··· CellN
Calculate single-cell ATAC counts per ACD by accumulating counts within each ACD
Single-cell ATAC counts matrix per ACD (M_ATAC)
ACD1
ACD2
ACD3
ACDM
Normalized to M̄_ATAC
Normalized single-cell ATAC counts matrix per ACD (N_ATAC)
Select the columns of N_ATAC with N̄_ATAC(j) ≥ M̄_ATAC to mitigate the influence of poorly sequenced cells
N_ATAC-Filtered
Cell1 ··· CellJ
Calculate Spearman correlation coefficient per ACD pair
Spearman correlation matrix (S_ATAC)
Calculate difference in Spearman correlation matrices
(ΔS_ATAC=S_ATAC,RAD21(-)-S_ATAC,Ctrl)
Differential Spearman correlation matrix (ΔS_ATAC)
ΔS_ATAC(i,j)
ACD index
B
Replicate #1
p<0.0001
Spearman correlation coefficient per ACD pair (S_ATAC(i,j))
auxin:
cis
trans
C
Replicate #2
p<0.0001
D
Ctrl
RAD21(-)
k_RAD21(-) = 3.70×10^-3
k_Ctrl = 5.06×10^-4
Normalized contact frequency per ACD pair

**Figure S2. Quantification of cross-domain chromatin co-accessibility.**

**(A)** The workflow for calculating differential Spearman correlation matrix ($\Delta S_{ATAC}$) per ACD pair before and after Cohesin depletion from single-cell ATAC-seq count outputs.

**(B-C)** Box plots of co-accessibility Spearman correlation coefficients ($S_{ATAC}(i,j)$) per ACD pair in *cis* (within the same chromosome) or *trans* (from different chromosomes) before and after Cohesin depletion for biological replicates #1 and #2. In the box charts, lower and upper whiskers represent 5%-95% values; the box represents the range from 25% to 75% percentile; the center line represents the median. Dotted line indicates the zero-change line. Non-parametric Wilcoxon test was used for statistical testing.

**(D)** Co-accessibility Spearman correlation coefficient ($S_{ATAC}(i,j)$) per ACD pair as a function of normalized Hi-C contact frequency per ACD pair before and after Cohesin depletion. $S_{ATAC}$ was plotted as a function of normalized contact frequency per ACD pair after a five-point smoothing. Shadow regions indicate standard error (S.E.) of the curves. The slope derived from linear regression of the curve for each condition was labelled below.

**A**

| Ectoderm lineage | Mesendoderm lineage | Chromosome | Distance (Mbps) |
|---|---|---|---|
| Gbx2 | Gpc1 | 1 | 2.92 |
| Foxj3 | Macf1 | 4 | 3.92 |
| Nf2 | Efemp1 | 11 | 24.08 |
| Nkx1-2 | Tead1 | 7 | 19.80 |
| Trim33 | Pitx2 | 3 | 25.89 |

**B**

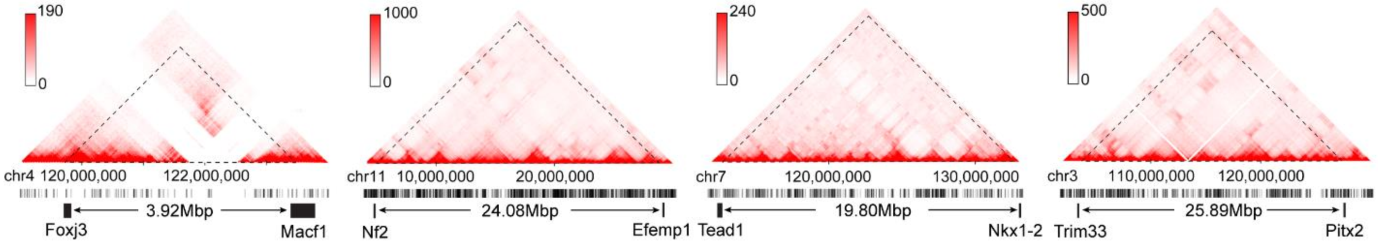


**C**

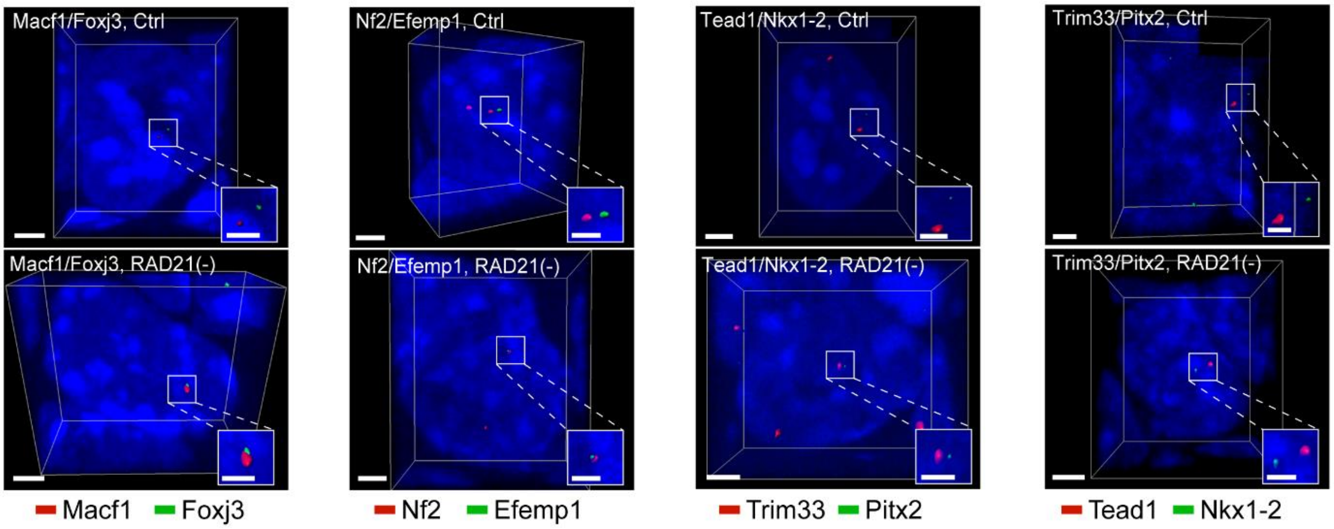

**Figure S3. Intron-FISH imaging of distant lineage-specific gene pairs *in cis.***

**(A)** The lineage and genomic location information for five pairs of developmental genes.

**(B)** Genomic positions of four pairs of representative lineage-specific genes with Hi-C and ATAC density information. The genomic distance between the two genes within each pair is labelled.

**(C)** Representative 3D *iso*-surface images of transcription bursting sites (intron-RNA-FISH) of gene pairs showed in (**A**) before and after Cohesin loss. Scale bar, 2μm. Inlet scale bar, 1μm.

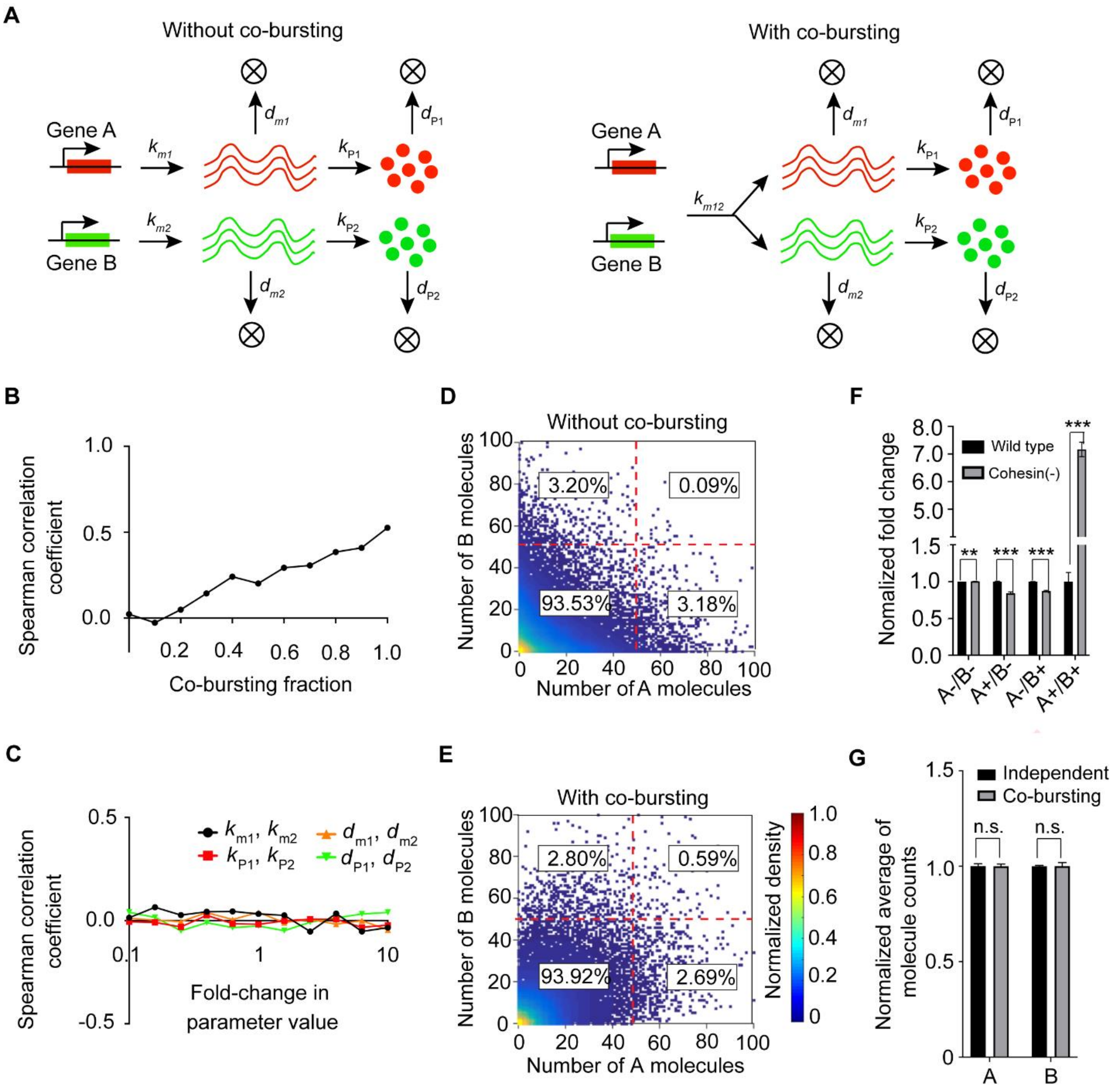
A
Without co-bursting
With co-bursting
Gene A
Gene B
B
Spearman correlation coefficient
Co-bursting fraction
C
Fold-change in parameter value
D
Without co-bursting
Number of B molecules
Number of A molecules
3.20%
0.09%
93.53%
3.18%
E
With co-bursting
2.80%
0.59%
93.92%
2.69%
Normalized density
F
Nomalized fold change
Wild type
Cohesin(-)
A-/B-
A+/B-
A-/B+
A+/B+
G
Independent
Co-bursting
Normalized average of molecule counts
n.s.

**Figure S4. Simulation by Gillespie algorithm predicts that increased gene co-activation has little effect on average gene expression.**

**(A)** Schematic diagrams illustrate gene co-expression simulation without or with co-bursting by Gillespie algorithm.

**(B)** Simulation results indicate that Spearman correlation coefficient increases along with the increase of co-bursting fraction for a pair of genes, A and B.

**(C)** Sensitivity analysis based on simulation with perturbed parameter values suggests that the change in Spearman correlation coefficient value is independent of other parameter values.

**(D-E)** Scatter plots of the numbers of A and B protein molecules without (**D**) or with (**E**) gene co-bursting. Each dot represents one cell. Density of the cell distribution was color-coded. Dotted lines divide the data plane into four quadrats, corresponding to four statuses of gene co-bursting: A-/B-, A+/B-, A-/B+ and A+/B+.

**(F)** Results from simulation based on Gillespie algorithm predict significant increase in the fraction of cells with co-activated lineage-specific gene pair (A and B) after Cohesin loss. Error bars indicate standard deviation (S.D.). Student's t-test was used for statistical testing of multiple independent stochastic simulations. *, $p < 0.05$; **, $p < 0.01$; ***, $p < 0.001$.

**(G)** Despite dramatic change in the fraction of cells with co-expressed genes (A+/B+), there is little change in average levels of A and B. The simulation was repeated for five times based on stochastic seeds and Student's t-test was used for statistical testing. Error bars indicate standard deviation (S.D.). n.s., not significant.

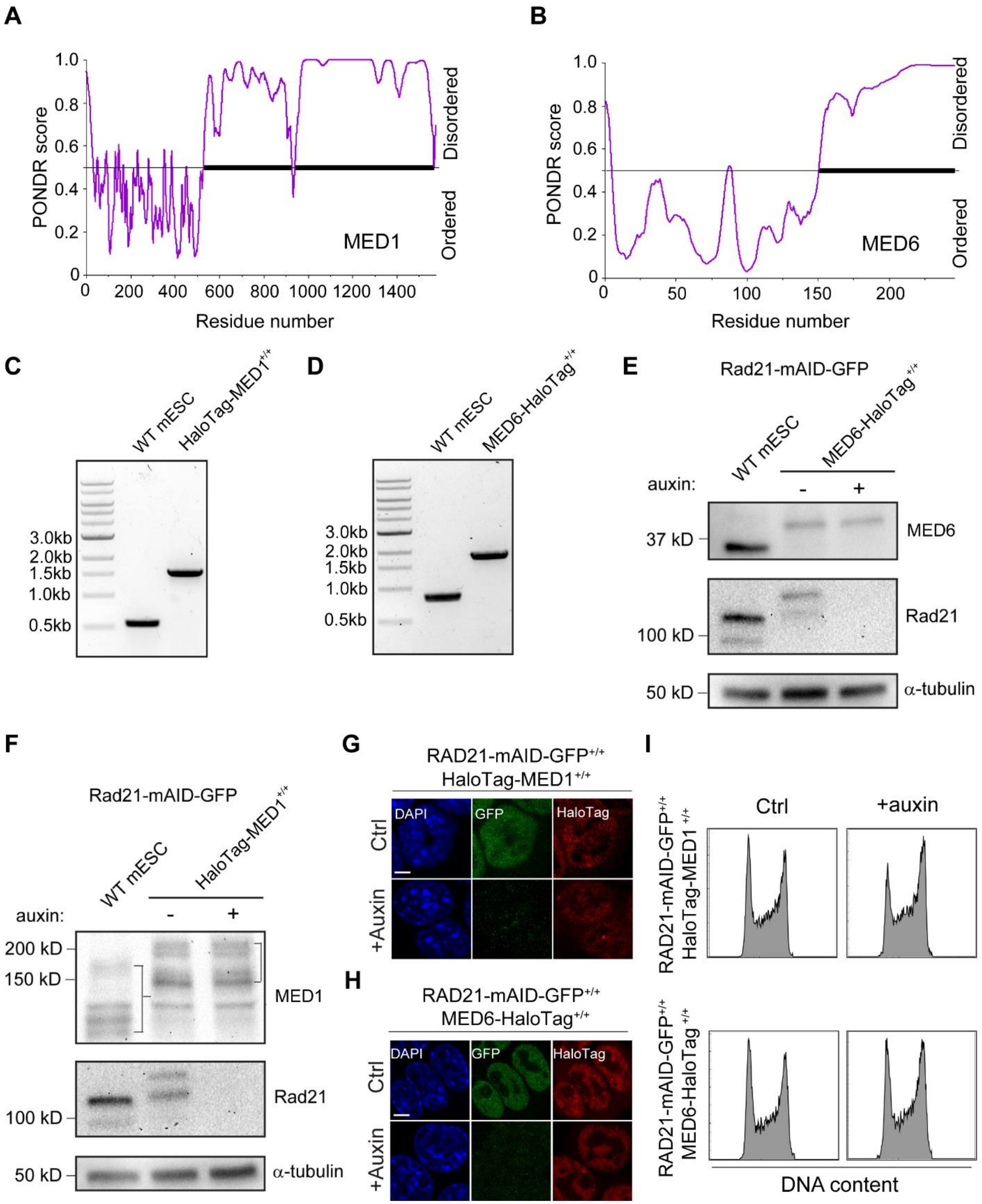
A
PONDR score
MED1
Disordered
Ordered
Residue number
B
PONDR score
MED6
Disordered
Ordered
Residue number
C
WT mESC
HaloTag-MED1+/+
3.0kb
2.0kb
1.5kb
1.0kb
0.5kb
D
WT mESC
MED6-HaloTag+/+
E
Rad21-mAID-GFP
MED6-HaloTag+/+
auxin:
37 kD
MED6
100 kD
Rad21
50 kD
α-tubulin
F
Rad21-mAID-GFP
HaloTag-MED1+/+
200 kD
150 kD
MED1
G
RAD21-mAID-GFP+/+
HaloTag-MED1+/+
DAPI
GFP
HaloTag
Ctrl
+Auxin
H
RAD21-mAID-GFP+/+
MED6-HaloTag+/+
I
Ctrl
+auxin
RAD21-mAID-GFP+/+
DNA content

**Figure S5. Construction of MED1/MED6-HaloTag knock-in cells.**

**(A-B)** PONDR score charts indicate predicted ordered and disordered regions within MED1 (**A**) and MED6 (**B**).

**(C-D)** PCR genotyping results showing bi-allelic fusion of HaloTag to MED1 (**C**; N-terminus) and MED6 (**D**; C-terminus)**.** Genomic DNA from wild-type mouse ES cells was used as the control.

**(E-F)** Western blots showing HaloTag-MED1 (**E**) and MED6-HaloTag (**F**) protein levels before and after RAD21 depletion by auxin-induced degron system. α-tubulin protein was blotted and used as a loading control.

**(G-H)** Fluorescence images showing RAD21-mAID-GFP (Green) and HaloTag-MED1 (**G**; Red) or MED-HaloTag (**H**; Red) levels without or with the auxin treatment (6 hrs). DNA was counter-stained with DAPI (Blue). Scale bar, 5μm.

**(I)** Flow-cytometry based cell cycle analysis of HaloTag-MED1 and MED6-HaloTag mouse ES cells before and after acute Cohesin loss (6 hrs).

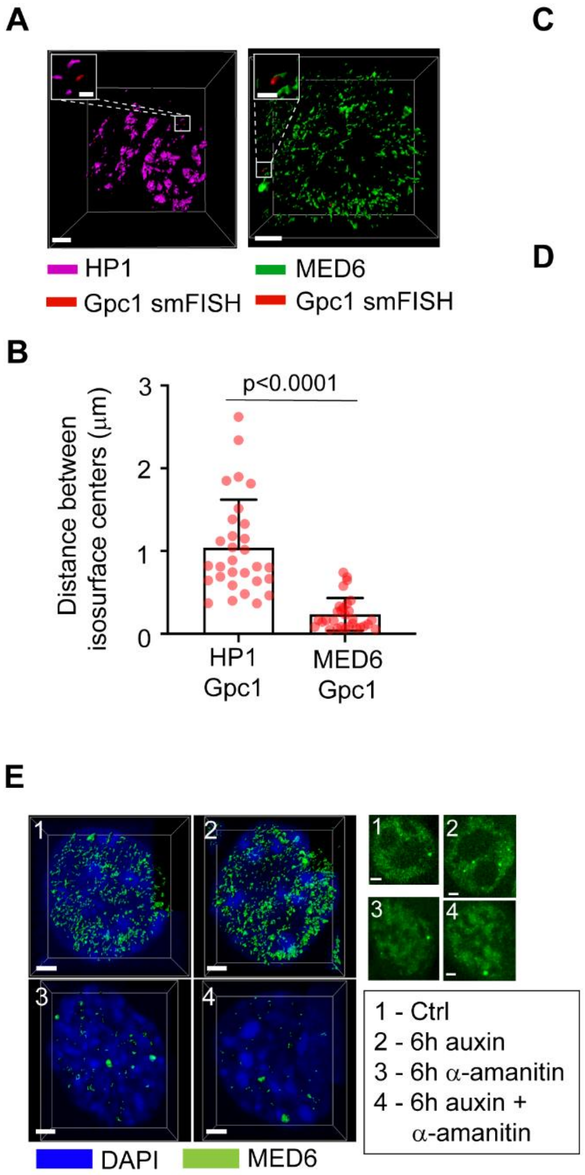

A
HP1
Gpc1 smFISH
MED6
Gpc1 smFISH
B
Distance between isosurface centers (μm)
p<0.0001
HP1 Gpc1
MED6 Gpc1
C
ATAC
MED1
Merge
#1
#2
D
Localization density
Pixel number
E
1 - Ctrl
2 - 6h auxin
3 - 6h α-amanitin
4 - 6h auxin + α-amanitin
DAPI
MED6
F
MED6 hub number
p=0.013
G
MED1 hub number
p=0.018

**Figure S6. Mediator hubs colocalize with ACDs.**

**(A)** 3D reconstruction shows the overlap between Gpc1 intron-FISH *iso*-surface and MED6-HaloTag hub *iso*-surface and the separation between Gpc1 intron-FISH *iso*-surface and HP1-GFP *iso*-surface. Scale bar, 2μm.

**(B)** Quantification of the physical distance between the centroid of Gpc1 intron-FISH signal and MED6 hub *iso*-surfaces and that between the centroid of Gpc1 intron-FISH intron-FISH singal and the nearest HP1 *iso*-surface. Non-parametric Wilcoxon test was used for statistical testing.

**(C)** One 2D section of two-color 3D ATAC and MED1 PALM images (Figure 4A) was used for spatial intensity correlation analysis in **C**. The original 3D localization maps were binned into 100 $nm^3$ cubic to generate 3D image volumes for both channels and one slice was selected for colocalization analysis. Scale bar, 2μm.

**(D)** One-dimensional intensity correlation analysis was performed for signals from two channels along selected line #1 and #2.

**(E)** Representative 3D reconstruction (left, 3D iso-surface rendering via Imaris) and raw images (upper right, one frame in Z-axis) of MED6 protein hubs in untreated, auxin treated (6hrs), α-amanitin treated (6hrs) and amanitin plus α-amanitin treated (6hrs) conditions. Cell nucleus was counter-stained with DAPI (Blue). Scale bar, 2μm.

(**F-G**) Quantification of MED6 (**F**) and MED1 (**G**) hub number in four different conditions as showed in (**E**). Mediator hubs from 10 cells were counted independently and Student's t-test was used for statistical testing.

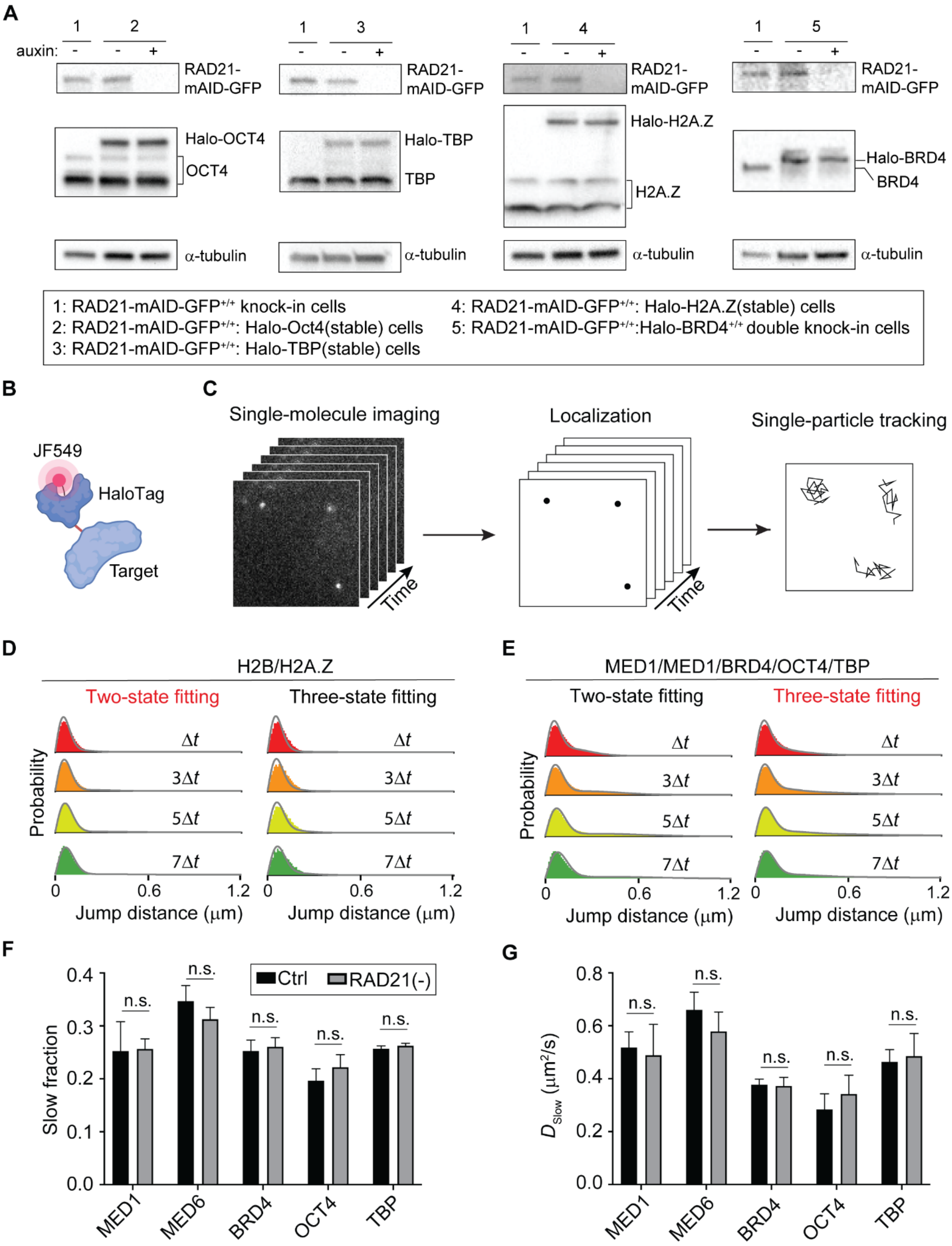
A
1
2
3
4
5
auxin:
RAD21-mAID-GFP
Halo-OCT4
OCT4
Halo-TBP
TBP
Halo-H2A.Z
H2A.Z
Halo-BRD4
BRD4
α-tubulin
1: RAD21-mAID-GFP+/+ knock-in cells
2: RAD21-mAID-GFP+/+: Halo-Oct4(stable) cells
3: RAD21-mAID-GFP+/+: Halo-TBP(stable) cells
4: RAD21-mAID-GFP+/+: Halo-H2A.Z(stable) cells
5: RAD21-mAID-GFP+/+:Halo-BRD4+/+ double knock-in cells
B
JF549
HaloTag
Target
C
Single-molecule imaging
Localization
Single-particle tracking
Time
D
H2B/H2A.Z
Two-state fitting
Three-state fitting
Probability
Δt
3Δt
5Δt
7Δt
Jump distance (μm)
E
MED1/MED1/BRD4/OCT4/TBP
F
Slow fraction
Ctrl
RAD21(-)
n.s.
MED1
MED6
BRD4
OCT4
TBP
G
D_Slow (μm²/s)

**Figure S7. Fast tracking reveals no apparent changes in Histone and TF dynamics after Cohesin loss.**

**(A)** Western blots indicate HaloTag fusion protein levels for either stably expressed (OCT4, TBP1 and H2A.Z) or endogenously labelled (BRD4) transcriptional regulators before and after RAD21 depletion. α-tubulin was used as the loading control.

**(B)** A diagram shows the labeling of HaloTag fusion proteins with JF549 dye.

**(C)** A schematic diagram illustrates the procedures for single-molecule imaging, localization, and tracking.

**(D-E)** Representative fittings of 2 state and 3 state model to jump histograms of H2B (**D**) and diverse transcriptional regulators (**E**) with variable $\Delta t$. Two state model that assuming that TFs alternates between one bound and one diffusive state, whereas three state model that assuming that TFs alternates between one bound, one slow and one fast diffusive state.

**(F)** Slow diffusive fractions for different transcriptional regulators before and after Cohesin removal calculated by Spot-On. The measurement was repeated for three times and Student's t-test was used for statistical testing. Error bars indicate standard deviation (S.D.). n.s., not significant.

**(G)** Diffusion coefficients for slow diffusive fractions as indicated in (**F**). The measurement was repeated for three times and Student's t-test was used for statistical testing. Error bars indicate standard deviation (S.D.). n.s., not significant.

### Supplementary Movies

#### Movie S1. Clustering of Chr2 intron-FISH puncta after Cohesin depletion.

The movie shows 3D *iso*-surface (magenta) reconstruction of Chr2 intron-FISH signals before and after Cohesin depletion (see also Figure 3B).

#### Movie S2. Mediator hubs overlap with accessible chromatin extensively.

Two-color 3D PALM imaging captures spatial distribution of both accessible chromatin sites (ATAC) (left) and MED1-HaloTag (middle) localizations (see also Figure 4A). As indicated, localization densities for both channels are color-coded.

#### Movie S3. Mediator hub fusion after Cohesin depletion.

The movie shows 3D *iso*-surface reconstruction of MED6 hubs before and after Cohesin depletion (see also Figure 4D). Individual hubs are color-coded by their volumes.

#### Movie S4. Time-lapse imaging of MED6 hubs upon Cohesin depletion.

The movie shows time-lapse imaging of MED6 hubs during acute Cohesion depletion (0~6 hrs) (see also Figure 4H). MED6 hubs are color-coded by their volumes.

### Supplementary Tables

#### Table S1. Genomic coordinates for accessible chromatin domains.

The table includes 776 identified accessible chromatin domains with their genomic locations in the mouse mm10 genome assembly.

#### Table S2. Chr2 intron-FISH gene list.

208 expressed genes in Chr2 were selected for intron-FISH experiment. The table includes gene ID, gene name, strand direction and their genomic locations in the mouse mm10 genome assembly.
